# High-throughput identification of RNA localization elements reveals a regulatory role for A/G rich sequences

**DOI:** 10.1101/2021.10.20.465152

**Authors:** Ankita Arora, Roberto Castro Gutierrez, Davide Eletto, Raquel Becker, Maya Brown, Andreas E. Moor, Holger A. Russ, J. Matthew Taliaferro

## Abstract

Hundreds of RNAs are enriched in the projections of neuronal cells. For the vast majority of them, though, the sequence elements within them that regulate their localization are unknown. To identify RNA elements capable of directing transcripts to neurites, we designed and deployed a massively parallel reporter assay that tested the localization regulatory ability of thousands of sequence fragments drawn from endogenous mouse 3′ UTRs. We identified peaks of regulatory activity within several 3′ UTRs and found that sequences derived from these peaks were both necessary and sufficient for RNA localization to neurites in mouse cells as well as active in human neurons. The active localization elements were enriched in adenosine and guanosine residues and were tens to hundreds of nucleotides long as shortening of the elements led to significantly reduced activity. Using RNA affinity and mass spectrometry, we found that the RNA-binding protein Unk was associated with the active localization elements. Depletion of Unk in cells reduced the neurite-enrichment of the elements, indicating a functional requirement for Unk in their trafficking. These results provide a framework for the unbiased, high-throughput identification of RNA elements and mechanisms that govern transcript localization.

## INTRODUCTION

In a variety of cell types across a range of species, thousands of RNA molecules are asymmetrically distributed within cells (Cajigas et al., 2012; Kislauskis et al., 1994; Lécuyer et al., 2007; Long et al., 1997). The localization of many of these RNAs is critical for specific cellular functions and developmental patterning (Ephrussi et al., 1991; Long et al., 1997; Nagaoka et al., 2012). In general, the localization of these RNAs is thought to be controlled by sequence elements, often termed “zipcodes”, that mark an RNA as one to be transported to a specific subcellular location (Hervé et al., 2004; Kislauskis et al., 1994; Meer et al., 2012). These sequences are often found in the 3′ UTR of the localized transcript and function through the recruitment of RNA-binding proteins (RBPs) that mediate the transport (Martin and Ephrussi, 2009). With the exception of a handful of examples (Engel et al., 2020), the identity of the localization regulatory sequence and the RBP that recognizes it are unknown.

Massively parallel reporter assays (MPRAs) have been used to identify regulatory mechanisms underlying a variety of gene expression regulatory processes including transcription (Arnold et al., 2013), RNA splicing (Mikl et al., 2019; Rosenberg et al., 2015), RNA stability (Rabani et al., 2017), lncRNA nuclear localization (Lubelsky and Ulitsky, 2018; Shukla et al., 2018), and protein abundance (Vainberg Slutskin et al., 2018). These methods test thousands of potential regulatory sequences in parallel in a single experiment. Sequences to be tested can either contain random sequence or naturally occurring sequence drawn from existing genomes. These sequences are integrated into reporter constructs, expressed in cells, and then populations of cells or nucleic acids are isolated based on the phenotype or process to be tested. The abundance of each individual member of the MPRA library within each cell or nucleic acid sample can then be quantified, and MPRA members associated with particular phenotypes or processes are identified. MPRAs therefore allow the rapid and unbiased detection of active regulatory elements from broad sequence pools. Despite the successful application of MPRAs to obtain insights into gene regulation, they have not been widely used in the study of RNA localization.

In recent years, transcriptomic approaches have been applied to the study of RNA localization, particularly in neuronal cells. In general, these approaches involve isolating and sequencing RNA from cell bodies and their projections (Cajigas et al., 2012; Middleton et al., 2019). Many groups have performed subcellular fractionations using cultured cells grown on microporous membranes that allow mechanical fractionation into cell body and neurite fractions (Ciolli Mattioli et al., 2019; Goering et al., 2020a; Taliaferro et al., 2016; Zappulo et al., 2017). High-throughput sequencing of RNA samples derived from these fractions have defined a fairly consistent set of transcripts that are reliably seen to be neurite-enriched (Goering et al., 2020a). However, mechanisms underlying how these transcripts become localized, including sequences within them required for localization, are almost completely unknown. In this study, we address this knowledge gap by employing a massively parallel reporter assay to assess the localization regulatory ability of sequences drawn from the 3′ UTRs of neurite-enriched transcripts.

## RESULTS

### Identification of 3′ UTRs sufficient for RNA localization activity

In order to study RNA localization in neuronal cells on a transcriptomic scale, we have developed and used a mechanical fractionation technique that makes use of cell growth on microporous membranes (Arora et al., 2021; Goering et al., 2020a; Hudish et al., 2020; Taliaferro et al., 2016) (**Figure 1A**). With this technique, cells are plated on the top of membranes whose pores allow neurite growth to the underside of the membrane. However, cell bodies are restricted to the top of the membrane. After cell growth and differentiation, the cells are mechanically fractionated into soma and neurite fractions by scraping the top of the membrane. RNA is collected from both fractions and can be analyzed by RT-qPCR or high-throughput sequencing.

**Figure 1.**
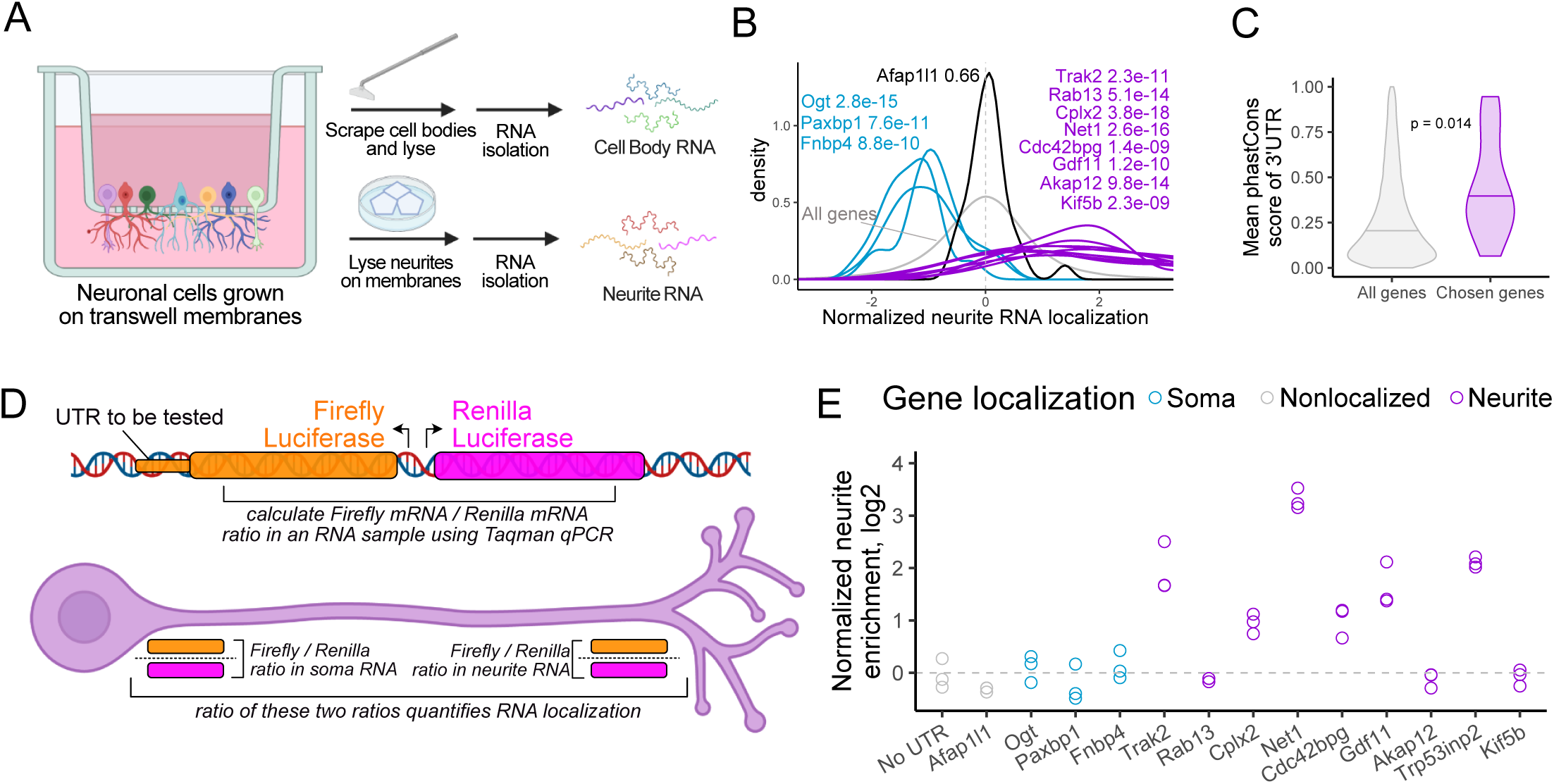
Identification of 3′ UTRs sufficient to drive RNA localization in neuronal cells. (A) Diagram of mechanical fractionation of neuron cells and analysis of subcellular transcriptomes. (B) Neurite-localized genes were identified through high-throughput RNA sequencing of compartment-specific transcriptomes from 32 mouse primary neuron and cell line-derived samples. Z-normalized neurite enrichments for the RNAs from selected genes are shown. Genes in purple were defined as repeatedly neurite-enriched. Genes in blue were defined as repeatedly soma-enriched, and the gene in black showed neither soma nor neurite enrichment. Wilcoxon p values represent the differences in neurite localization distributions between the indicated genes and all genes (gray). (C) PhastCons conservation scores for the 3′ UTRs of all genes (gray) and the chosen neurite-enriched genes (purple). (D) Diagram of RT-qPCR experiment. Reporter plasmids expressing the firefly and renilla luciferase transcripts are integrated into the genome through Cre-mediated recombination and are expressed from a bidirectional promoter. Sequences whose RNA localization activity will be tested are fused onto the 3′ UTR of Firefly luciferase. The ratio of firefly to renilla luciferase transcripts in soma and neurite samples is measured using Taqman qPCR. Comparing these ratios in soma and neurite samples quantifies localization of the firefly luciferase transcript. (E) 3′ UTRs of the indicated genes were fused to firefly luciferase and the neurite localization of the resulting transcript was quantified using RT-qPCR. The neurite localization of the firefly luciferase with no added 3′ UTR was used as a control.

We have performed this technique on 32 mouse samples ranging from neuronal cell lines to primary cortical neurons (Goering et al., 2020a; Hudish et al., 2020; Taliaferro et al., 2016). By amalgamating these results, we compiled repeated observations of the RNA localization patterns of thousands of transcripts and identified those which were reproducibly enriched in neurites. We chose to focus on 9 genes whose transcripts were strongly neurite-enriched (**Figure 1B, S1A**) with the goal of identifying sequences within these transcripts that regulate their localization. As controls, we also chose three genes that were reproducibly soma-enriched and one that was neither soma-nor neurite-enriched (**Figure 1B, S1A**).

To narrow the possible sequence search space, we focused on the 3′ UTR sequence of each of these genes as 3′ UTRs have been found to regulate the localization of many RNAs (Goering et al., 2020b; Meer et al., 2012; Taliaferro et al., 2016; Tushev et al., 2018). We found that the 3′ UTRs of the chosen neurite-localized genes were more conserved than other 3′ UTRs, suggesting that they may contain functional localization elements (**Figure 1C**). Using RNAseq data from fractionated human motor neurons (Goering et al., 2020a; Hudish et al., 2020), we found that the localization of the human orthologs of these transcripts was very similar to their mouse counterparts, further suggesting that conserved elements within these transcripts mediate their transport (**Figure S1B**).

To directly test if the 3′ UTRs of these transcripts contained RNA localization regulatory elements, we incorporated their 3′ UTRs into a reporter system we designed that contained firefly and renilla luciferases driven by a bidirectional doxycycline-sensitive promoter. The 3′ UTRs to be tested were fused to the firefly luciferase transcript while the 3′ UTR of the renilla luciferase transcript was kept constant, allowing it to serve as an internal control. We then site-specifically integrated this construct into the genome of mouse CAD cells through cre/loxP-mediated recombination (Khandelia et al., 2011). Using single molecule fluorescence in situ hybridization (smFISH) probes against the firefly transcript, we found that the 3′ UTRs of some neurite-localized genes were sufficient to localize a reporter transcript to neurites (**Figure S1C**).

We then fractionated these reporter-containing cells using mechanical separation into soma and neurite fractions (**Figure 1A**) and calculated the relative firefly luciferase and renilla luciferase transcript levels in the resulting RNA samples using Taqman RT-qPCR. By comparing these relative firefly to renilla luciferase transcript ratios in the soma and neurite fractions (i.e. calculating a ratio of ratios), we quantified the localization of various firefly luciferase fusion constructs (**Figure 1D**).

As a control, we used a construct in which no endogenous 3′ UTR was fused to the firefly luciferase transcript. We set the resulting neurite to soma ratio of firefly to renilla ratios of this construct to one. We then tested the effect of fusing the 3′ UTRs of the identified neurite- and soma-enriched genes on this localization-quantifying ratio of ratios (**Figure 1E**). We found that fusion of the 3′ UTR of 6 of the 9 neurite-localized genes to firefly luciferase resulted in substantially increased neurite enrichment of the firefly luciferase transcript. Conversely, none of the four control 3′ UTRs had an effect on the neurite enrichment of firefly luciferase RNA. We therefore concluded that RNA localization regulatory elements were located somewhere within the identified active 3′ UTRs.

### Design of a massively parallel reporter assay

In order to define RNA localization regulatory elements within active 3′ UTRs, we designed an MPRA covering the 3′ UTRs of chosen genes. Because previously identified localization regulatory elements were quite large (50-250 nt) (Chabanon et al., 2004; Engel et al., 2020), we chose to use long oligonucleotides that were densely tiled in order to identify active elements with high resolution. We designed 260 nt oligonucleotides that tiled the chosen 3′ UTRs with one oligonucleotide every 4 nt (**Figure 2A**). This resulted in an oligonucleotide library with approximately 8100 members in which each individual nucleotide within a UTR was incorporated into 65 distinct oligonucleotides (**Supplementary files 1 and 2**). As positive controls, we included 3 oligonucleotides that covered the entirety of the 150 nt neurite-localized long noncoding RNA BC1 (Brosius and Tiedge, 2001). As negative controls, we included oligonucleotides that tiled the 3′ UTRs of four RNAs that were not neurite-enriched (**Figure 1E**) as well as oligonucleotides that tiled the length of the nucleus-restricted lncRNA Malat1.

**Figure 2.**
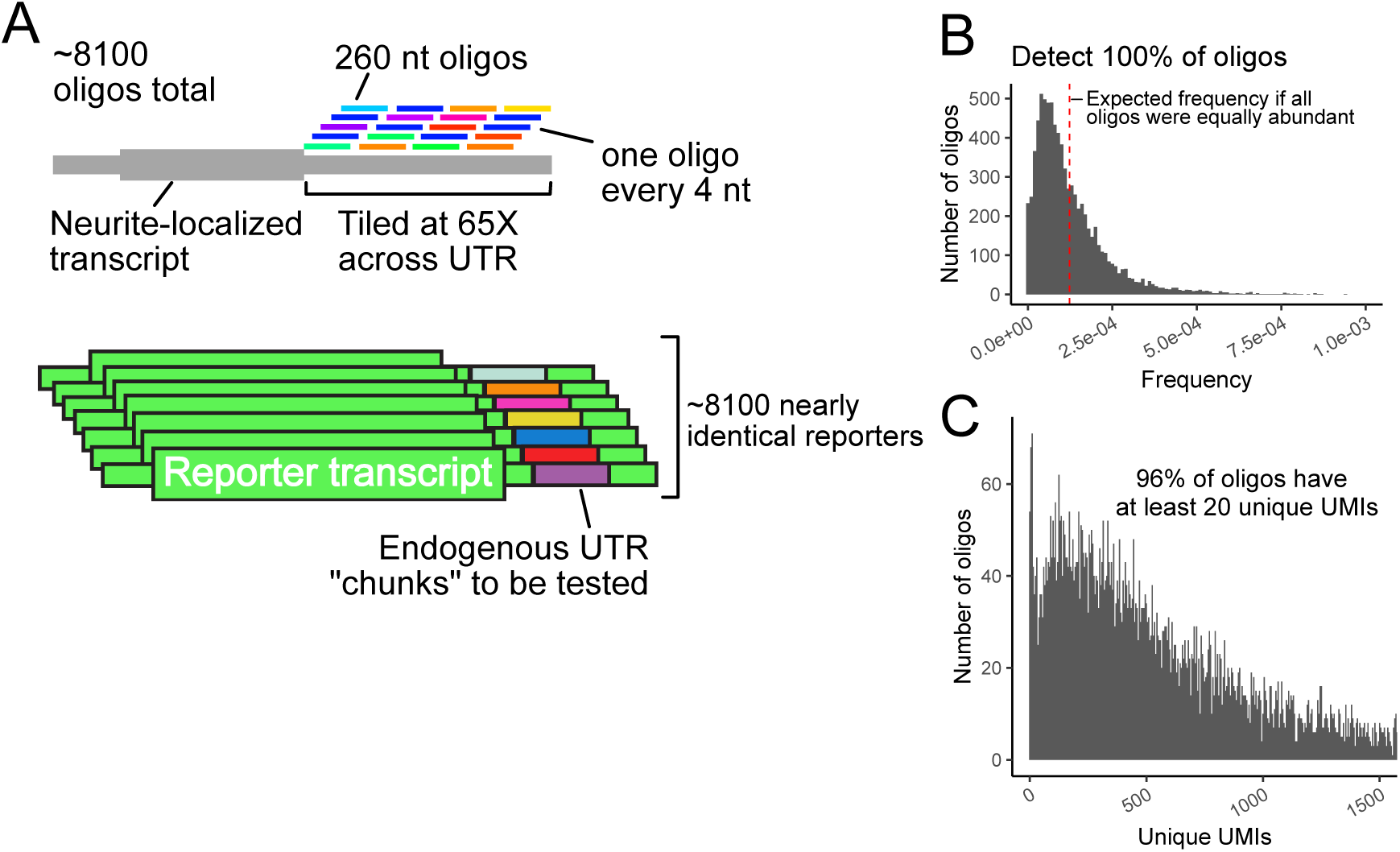
Design of MPRA and QC of synthesized oligonucleotide pool. (A) Oligonucleotides of length 260 nt were designed against the 3′ UTRs of chosen genes. Neighboring oligonucleotides were spaced 4 nt from each other, giving an average coverage of 65X per nucleotide. These oligonucleotides were then integrated into the 3′ UTR of reporter transcripts, generating a library of reporters. (B) Distribution of oligonucleotide abundances in the synthesized pool. (C) Distribution of oligonucleotide abundances in the integrated GFP reporter transcript in CAD neuronal cells.

We were concerned that high-throughput sequencing read aligners like Bowtie2 (Langmead and Salzberg, 2012) might have difficulty correctly assigning reads to oligonucleotides that differ from each other by only 4 nt steps. To assess this, we created multiple simulated MPRA libraries of ten thousand 260 nt UTR fragments. In these libraries, oligonucleotides were separated by 2, 5, or 10 nt. We created 10 million mock sequencing reads from these libraries, including mimicked oligonucleotide synthesis and sequencing errors (Pfeiffer et al., 2018). We found that decreasing the step size between oligonucleotides resulted in more reads being incorrectly assigned (**Figure S2A**). We observed that the incorrectly assigned reads had lower mapping qualities (**Figure S2B**). We therefore reasoned that allowing Bowtie2 more chances to find optimal alignments (controlled by the parameter -D) might increase performance. Increasing the value of the parameter from its default of 15 to 150 allowed Bowtie2 to correctly assign 100% of reads from the 2 nt step library (**Figure S2C**). We therefore concluded that with the modified parameter, Bowtie2 can accurately assign reads from high density MPRA libraries.

### Quality control of MPRA reagents and procedure

To check the quality of the oligonucleotide library, we analyzed it using paired-end high-throughput sequencing. We detected 100% of the expected oligonucleotides in the library and found that most of them were approximately equally abundant (**Figure 2B**). We found that approximately 65% of the oligonucleotides contained no mutations, insertions, deletions (**Figure S2D**). However, the majority of indels and mutations occurred in regions of the oligonucleotide sequenced by only one read of the paired-end reaction (**Figure S2E**). In the middle of the oligonucleotide where the reads overlapped, we required that both reads contain a mutation in order to call a mutation. Because the observed mutation rate is much lower in the paired end overlap, we conclude that the majority of observed errors are sequencing errors, and that the oligonucleotide library is largely error-free.

The oligonucleotide library was cloned into two places in a single plasmid: once into the 3′ UTR of a gene encoding GFP and once into the 3′ UTR of a gene encoding firefly luciferase. Each plasmid molecule therefore contained two different oligonucleotides incorporated into two different doxycycline-inducible reporter transcripts.

To express this library of reporter transcripts at moderate and controllable levels, we site-specifically integrated constructs into the genomes of CAD and N2A mouse neuronal cells using cre/loxP-mediated recombination (Khandelia et al., 2011). Using this strategy ensured that each cell is only expressing one firefly luciferase reporter RNA and one GFP reporter RNA out of the 8100 possible. Further, each cell expresses its reporters from the same genomic locus. We reasoned that this would avoid the overwhelming of the RNA processing and transport machinery that may occur with the high expression typical of transient transfection.

In order to use this strategy, we needed to know how many independent integration events we could generate. To calculate this, we integrated a plasmid containing a randomized 15 nt segment into 6 million CAD and N2A cells. After selecting for integrants with puromycin, we used targeted high-throughput sequencing to determine the number of unique 15 nt segments that were genomically integrated. We observed approximately 600,000 unique CAD integration events and approximately 200,000 unique N2A integration events (**Figure S2F**), giving an cre-mediated integration efficiency of 5-10%, which agreed well with previously published values (Lanza et al., 2012). We therefore concluded that integrating our 8100 reporter constructs was feasible.

After integrating our MPRA library, we interrogated the complexity of the integrated oligonucleotides using targeted RNA sequencing. We detected 99% of the oligonucleotides in the library incorporated into the GFP reporter site in CAD cells. 96% of the oligonucleotides could be quantified with high confidence, as evidenced by them being associated with at least 20 unique molecular identifiers (UMIs) incorporated during reverse transcription (**Figure 2C**). Similarly, we detected 97% of the oligonucleotides in the GFP reporter in N2A cells, with 86% of them being reliably quantified (**Figure S2F**).

### MPRA results are reproducible across cell lines and reporter transcript scaffolds

We fractionated CAD and N2A cells containing the MPRA library into cell body and neurite fractions in quadruplicate. Using targeted RNAseq, we then quantified the relative abundance of each oligonucleotide in the firefly luciferase and GFP reporters in each fraction (**Tables S1-S4**). UMIs that were incorporated during reverse transcription were used for quantification to exclude PCR amplification artifacts.

We found that oligonucleotide abundances within samples clustered by cellular compartment and that abundances within compartments were highly similar (**Figure 3A, S3A**). We then identified oligonucleotides that were significantly enriched in a compartment using DESeq2 (Love et al., 2014), using a False Discovery Rate (FDR) cutoff of 0.01. Using these parameters, we identified 379 and 220 oligonucleotides that were neurite- and soma-enriched, respectively, in the GFP reporter construct in CAD cells. 269 and 132 oligonucleotides were neurite- and soma-enriched, respectively, using the GFP construct in N2A cells.

**Figure 3.**
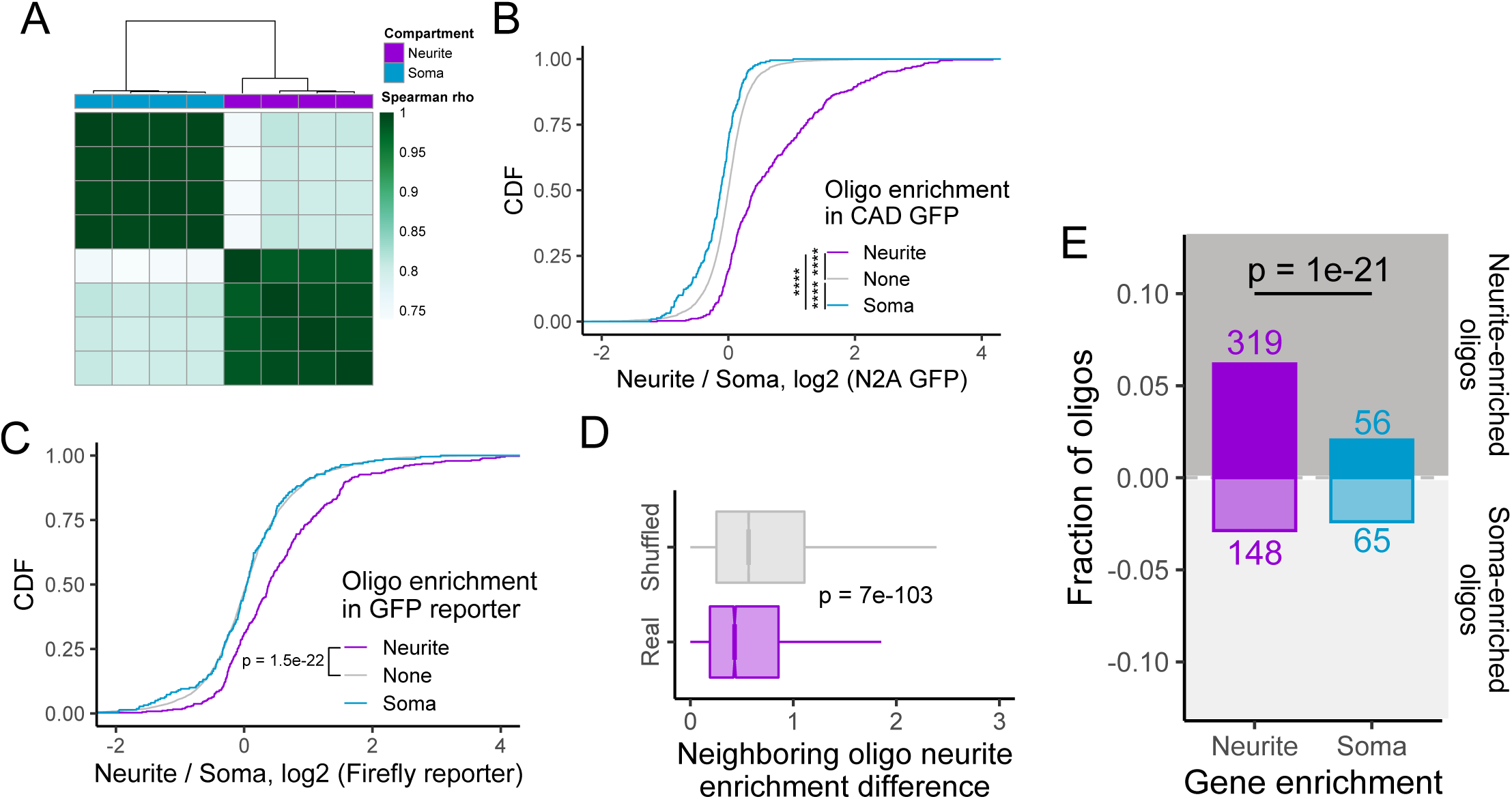
Overview and QC of MPRA results. (A) Hierarchical clustering of oligonucleotide abundances from the GFP reporter in CAD cells. (B) Concordance of results across cell lines. Neurite enrichment in N2A GFP samples of oligonucleotides defined as soma-(blue), neurite-(purple), or non-localized (gray) in CAD GFP samples. (C) Concordance of results across reporter scaffolds. Neurite enrichment in CAD firefly luciferase samples of oligonucleotides defined as soma-, neurite-, or non-localized in CAD GFP samples. (D) Distribution of absolute differences in neurite enrichment values for neighboring oligonucleotides. As a control, the positional relationship between all oligonucleotides was randomly shuffled, and the difference in neurite enrichment for neighboring oligonucleotides were recalculated. (E) Number of significantly neurite- and soma-enriched oligonucleotides among those drawn from UTRs from neurite- and soma-enriched genes. All significance tests were performed using a Wilcoxon rank-sum test. p value notation: * < 0.05, ** < 0.01, *** < 0.001, **** < 0.0001.

To determine if oligonucleotide enrichments were consistent across cell lines, we compared the neurite-enrichments in N2A cells of oligonucleotides defined as neurite-, soma- or non-enriched in CAD cells. We found that oligonucleotides that were neurite-enriched in CAD cells had significantly higher neurite enrichments in N2A cells than non-localized oligonucleotides. Conversely, oligonucleotides that were soma-enriched in CAD cells had significantly lower neurite enrichments than non-localized oligonucleotides (**Figure 3B**). Further, neurite enrichments for all oligonucleotides in CAD and N2A cells were significantly correlated with each other (**Figure S3B**,**C**).

To determine if oligonucleotide enrichments were consistent across reporter constructs, we similarly compared neurite enrichments for oligonucleotides embedded in the GFP and firefly luciferase reporters. Oligonucleotides that were significantly neurite-enriched when embedded within the GFP reporter had significantly higher neurite enrichments than expected when embedded in the firefly luciferase reporter (**Figure 3C**), indicating a broad concordance of results between reporter transcripts. This suggests that the localization observed localization activity is independent of the broader sequence context of the reporter.

If the observed data were reliable, then we would expect that two oligonucleotides who neighbor each other in their position within a UTR to have similar neurite enrichments since they share 252 nt of sequence. To assess this, we calculated absolute differences in neurite enrichments for all neighboring oligos. As a control, we shuffled the positions of oligonucleotides along UTRs but within a gene and repeated the analysis. We observed that the neurite enrichments of neighboring oligonucleotides were significantly more similar to each other than in the shuffled control (**Figure 3D**). We then performed this analysis for each gene individually. Interestingly, in the neurite-enriched genes, neighboring oligonucleotides had similar neurite enrichments. However, this was not true for the soma-enriched genes (**Figure S3F**).

### Oligonucleotides drawn from neurite-enriched genes are more likely to be neurite-enriched than those drawn from soma-enriched genes

If our results were to make biological sense, we would expect that oligonucleotides drawn from neurite-enriched genes would be more likely to be neurite-enriched than those drawn from soma-enriched genes. Reassuringly, we found that oligonucleotides from neurite-enriched genes were 3 times as likely to themselves be neurite-enriched than those contained within soma-enriched genes (p = 1e-21, binomial test) (**Figure 3E**). When we considered each gene individually, most neurite-enriched genes contained far more neurite-enriched oligos than soma-enriched oligos. This includes our positive control, BC1, for whom two out of its three constituent oligos were significantly neurite-enriched (**Figure S3G**).

### Neurite-enriched oligonucleotides cluster together along the 3′ UTR to define peaks of activity

We then arranged oligonucleotides according to their position along their parental 3′ UTRs and plotted their neurite enrichments. Encouragingly, we found that for all of the 3′ UTRs that were sufficient to direct localization to neurites, peaks of activity within the 3′ UTR were clearly visible (**Figure 4A-E, S4A**). These peaks represent active RNA elements within the 3′ UTR that regulate RNA localization. We were not able to detect peaks of activity in the 3′ UTRs of genes that were not neurite-localized (**Figure 4F, S4B**), further demonstrating the specificity of the experiment. Intriguingly, we did detect one clear peak of activity within the body of Malat1 (**Figure S4B**). Importantly, these peaks span dozens of oligonucleotides, meaning that fully independent yet related oligonucleotides behaved similarly in the MPRA, giving us confidence in the results.

**Figure 4.**
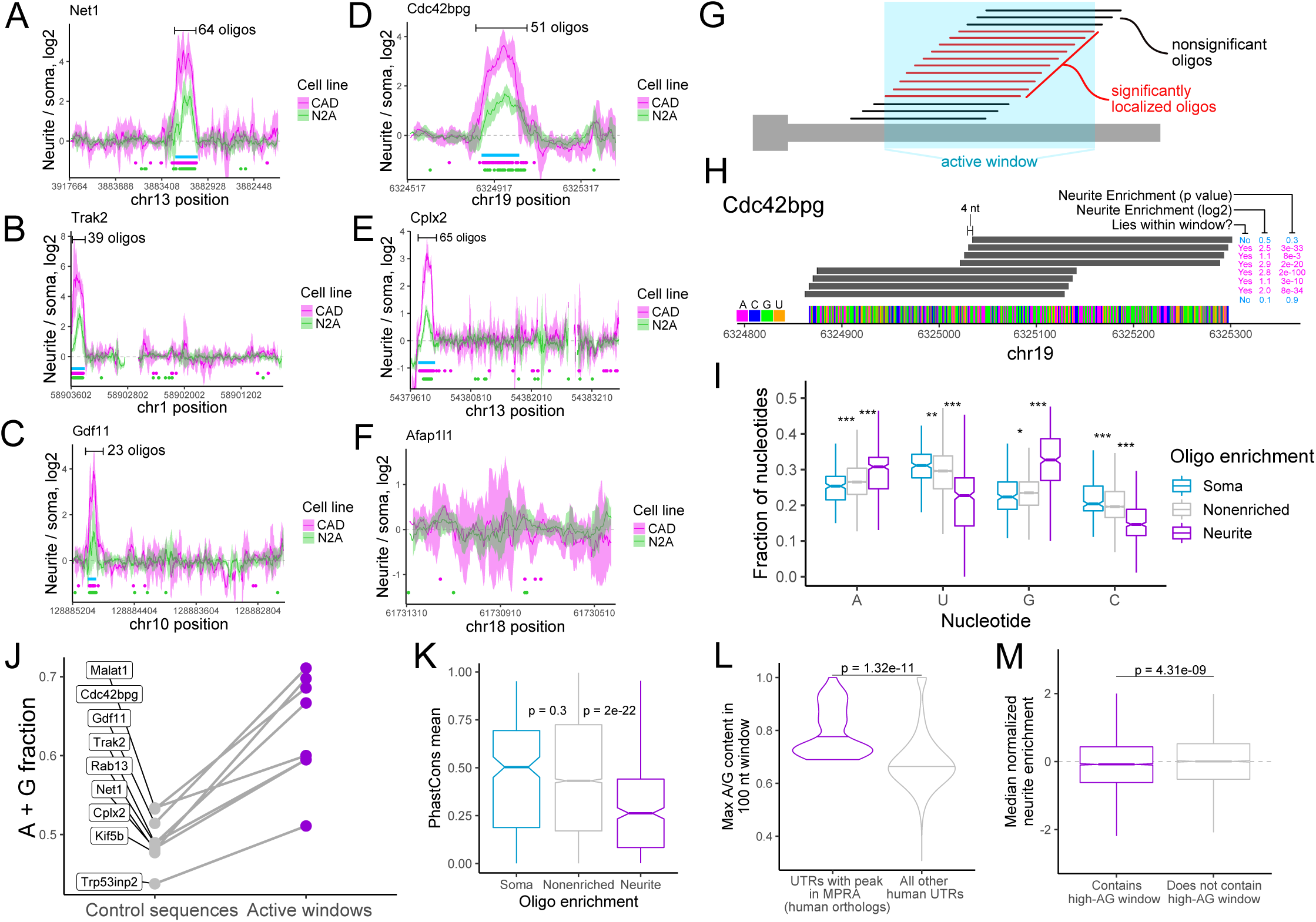
Identification of UTR regions with RNA localization activity and their properties. (A-F) Neurite enrichments of oligonucleotides as a function of their location within the gene’s UTR. These plots are using GFP reporter data from CAD (pink) and N2A (green) cells. Lines represent a sliding average of 8 oligonucleotides, and the ribbon represents the standard deviation of neurite enrichment for the oligonucleotides in the sliding window. Dots below the lines represent the locations of significantly neurite-localized oligonucleotides (FDR < 0.05). Blue boxes represent the locations of “active windows” defined using the CAD data. Note that the 3′ UTR of Afap1l1 was not expected to contain active oligonucleotides. (G) Definition of active window sequences. (H) Resolution of active sequences afforded by high density oligo design. Neighboring oligos show vast differences in activity even though they lie only 4 nt apart. (I) Nucleotide content of oligos defined as significantly soma-, neurite, and non-localized using CAD GFP data. (J) A/G content of active windows and inactive sequences in the indicated UTRs. (K) Conservation scores of soma-, neurite-, and non-localized oligonucleotide sequences. (L) Maximum A/G content of 100 nt windows for the 3′ UTRs of the human orthologs of the genes that contain active peaks in the MPRA. (M) Median Z-normalized neurite enrichment across 32 RNA localization experiments for genes whose 3′ UTR contains a 100 nt window with at least 70% A/G content (purple) or does not contain a high A/G content window (gray). All significance tests were performed using a Wilcoxon rank-sum test. p value notation: * < 0.05, ** < 0.01, *** < 0.001, **** < 0.0001.

From this data, we formalized an approach for defining windows of active oligonucleotides (see Methods). Essentially, contiguous stretches of significantly neurite-enriched oligos (FDR < 0.01) were combined to define “active windows” (**Figure 4G**). The 4 nt tiling distance between neighboring oligonucleotides affords a high degree of resolution of the boundaries of these windows. Two oligonucleotides, although separated by only 4 nt, can show dramatically different neurite enrichments (**Figure 4H**).

### Characteristics of neurite-enriched oligonucleotides

We then asked if neurite-enriched oligonucleotides were enriched for specific properties that might define or contribute to their activity. We found that neurite-enriched oligonucleotides had significantly higher adenosine and guanosine contents than nonlocalized oligonucleotides (**Figure 4I**) and were also enriched for A/G-rich kmers (Rasmussen et al., 2013) (**Figure S4C**). Conversely, neurite-enriched oligos were depleted for cytosine (**Figure 4I**), consistent with recent reports that C-rich elements are important for nuclear retention of RNA molecules (Lubelsky and Ulitsky, 2018; Shukla et al., 2018). Further, all active window sequences had higher average A/G contents than the rest of the 3′ UTR that contained them (**Figure 4J, S4D-F**). These results suggest that A/G richness is a key contributor to the ability of an RNA element to direct transcript localization.

Multiple previous reports have emphasized a likely role for RNA secondary structure in the definition of RNA localization regulatory elements (Hamilton and Davis, 2011; Hamilton et al., 2009). To assess the secondary structure character of our oligonucleotides, we computationally folded them using RNAfold (Lorenz et al., 2011). Surprisingly, we found no difference in the minimum free energy of soma-, non-, and neurite-enriched oligonucleotides (**Figure S4G**). We did, however, find that neurite-enriched oligonucleotides were significantly more likely to contain predicted G-quadruplexes (**Figure S4H**), consistent with previous reports of the ability of G-quadruplex RNA sequences to drive RNA localization to neurites (Goering et al., 2020a; Subramanian et al., 2011).

If the identified sequences were functional, we would expect them to be conserved. Unexpectedly, the sequences of neurite-enriched oligos were less conserved than those of soma- or non-enriched oligonucleotides (**Figure 4K**). We suspect that this may be due to two factors. First, if a defining characteristic of our identified active elements is their A/G richness, the exact nucleotide sequence of the element may be less important and therefore less likely to be conserved. Second, it may not be necessary for these elements to be *positionally* conserved and therefore alignable across genomes. It may be that the mere *presence* of the element anywhere within the 3′ UTR is sufficient for function. If this were true, then the 3′ UTRs of the human orthologs of the genes with active windows should also contain stretches with high A/G content. We found that this was, in fact, the case (**Figure 4L**), suggesting that similar mechanisms regulate the localization of human and mouse orthologous transcripts.

However, the presence of an A/G-rich window on its own does not seem sufficient to predict whether or not a transcript will be localized. We binned genes into those that contained a high A/G window in their 3′ UTR (a 100 nt stretch of at least 70% A/G) and those that do not. By comparing dozens of subcellular RNAseq datasets from a variety of neuronal cell types (Goering et al., 2020a; Minis et al., 2013; Taliaferro et al., 2016; Tushev et al., 2018; Zappulo et al., 2017), we found that, on average, genes with high A/G windows in their 3′ UTR were actually slightly less neurite-enriched than those without high A/G windows (**Figure 4M**). This suggests that additional information beyond an A/G rich window is required to form an active localization regulatory element.

### Identified oligonucleotides are necessary and sufficient for RNA transport

To verify the ability of individual oligonucleotides to direct RNA transport, we created reporter transcripts containing oligonucleotides identified as active by the MPRA. For each MPRA-defined peak of activity along a 3′ UTR, we defined a “peak” oligo that lay near the center of the peak of activity (**Figure 5A**). We selected peak oligonucleotides from two genes, *Net1* and *Trak2*, fused them to our reporter construct, and assayed the ability of the peak oligonucleotides to drive neurite enrichment of the transcript by smFISH. We found that in contrast to a reporter transcript lacking an oligonucleotide fusion, reporters containing peak oligonucleotides were highly enriched in neurites (**Figure 5B**), demonstrating that the peak oligonucleotides were sufficient for RNA transport.

**Figure 5.**
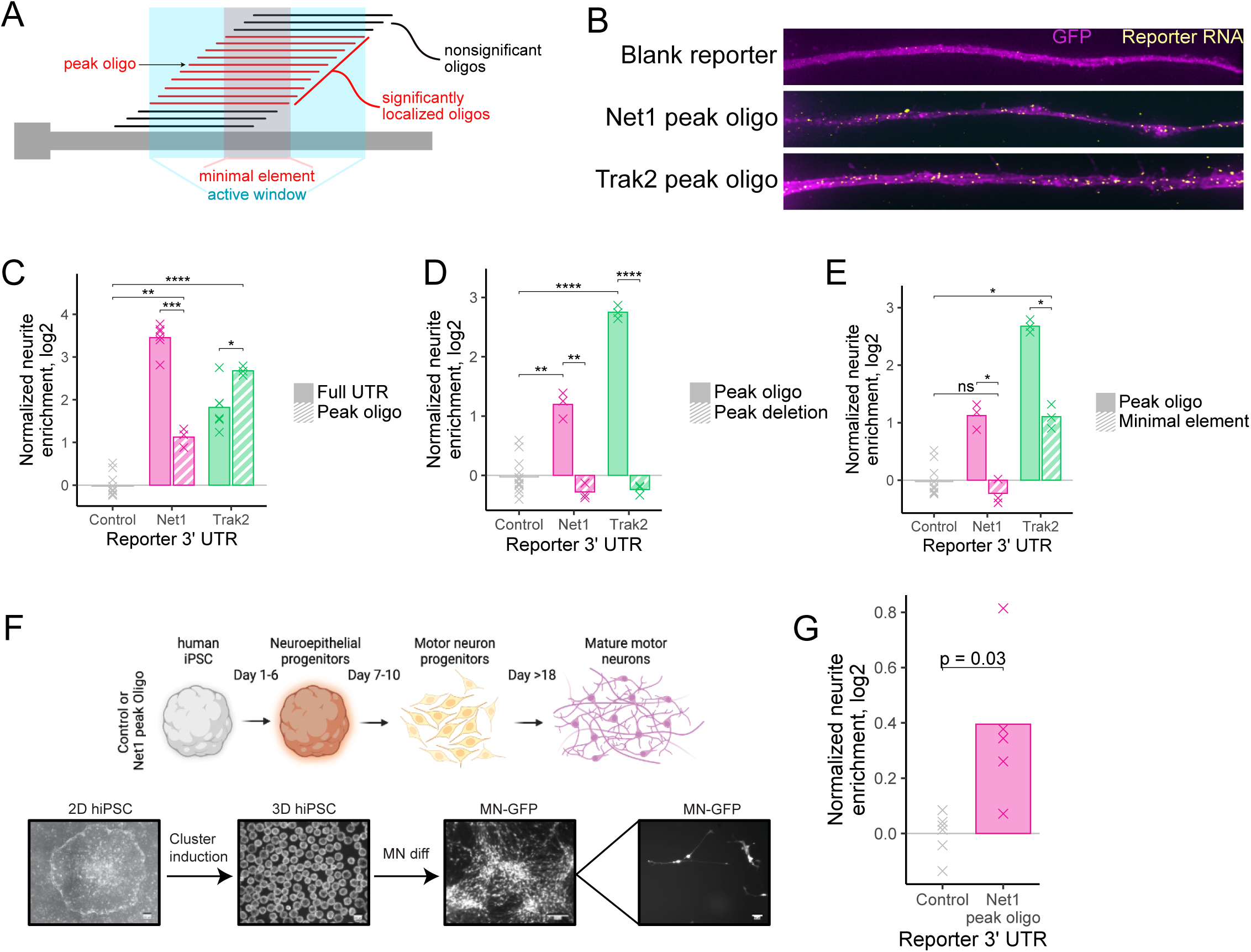
Experimental validation of active sequences. (A) Active sequence windows (blue) were defined as the sequences present within any oligonucleotide contained within a stretch of neurite-localized oligonucleotides. Minimal element sequences were defined as those present within all oligonucleotides of a stretch of neurite-localized oligonucleotides. Peak oligonucleotides were those with high localization activity, often found at the center of an active window. (B) smFISH of reporter constructs containing peak oligonucleotide sequences. (C) RNA localization activity, as assayed by RT-qPCR, of peak oligonucleotide sequences and the full UTRs from which they were drawn. Observed neurite enrichments for each construct were compared to the enrichment of a control reporter construct lacking an active 3′ UTR. (D) As in C, RNA localization activity of reporter constructs containing peak oligonucleotide or full UTRs lacking the sequences of the peak oligonucleotides. (E) RNA localization activity of reporter constructs containing peak oligonucleotides or minimal elements as defined in A. (F) Schematic of differentiation of human iPS cells containing reporter constructs into motor neurons. (G) RNA localization activity, as assayed in human motor neurons, of a reporter construct containing the peak oligonucleotide from the mouse Net1 gene. All significance tests were performed using a t-test. p value notation: * < 0.05, ** < 0.01, *** < 0.001, **** < 0.0001.

We then compared the RNA localization of reporters containing either peak oligonucleotides or the entire 3′ UTR from which the peak oligonucleotide was obtained using cell fractionation and RT-qPCR (**Figure 1D**). We found that for *Net1*, although the peak oligonucleotide contained activity, it could not drive neurite-enrichment to the same extent as its parental 3′ UTR. Conversely, for *Trak2*, the peak oligonucleotide was slightly more active than its parental 3′ UTR (**Figure 5C**). These results suggest that contextual effects of the sequence surrounding the peak oligonucleotide may influence its regulatory ability.

Next, we compared the localization activities of 3′ UTRs either containing or specifically lacking the identified peak oligonucleotides. For both *Net1* and *Trak2*, we found that removal of the peak oligonucleotide completely abolished the activity contained within the 3′ UTR (**Figure 5D**), indicating that the peak oligonucleotide is necessary for localization of the reporter transcripts.

We then attempted to identify active elements with higher resolution by defining “minimal elements”. We reasoned that the sequence elements that were driving RNA localization were likely to be those in common to all oligonucleotides contained within an active window (**Figure 5A**). Essentially, while an active window would contain the union of all active oligonucleotides within a given region, a minimal element would contain their intersection. We defined minimal elements for all identified peaks of activity with the lengths of the minimal elements ranging from 56 nt to 224 nt (**Figure S5A**). The minimal elements for *Net1* and *Trak2* were 56 and 107 nt, respectively.

For both *Net1* and *Trak2*, reporters containing minimal elements were significantly less localized than reporters containing peak oligos, despite the fact that the minimal elements are core subsequences of the peak oligonucleotides (**Figure 5E**). Peak oligonucleotides of length 260 nt contained significantly more activity than their constitutive minimal elements of length 56 and 107 nt, suggesting that RNA localization elements are generally much larger than regulatory elements that control other RNA metabolic processes. This is in line with the sizes of previously defined localization regulatory elements (Engel et al., 2020; Hervé et al., 2004).

### Peak oligonucleotides are sufficient to drive RNA localization in human motor neurons

To assess the ability of the peak oligonucleotides to regulate RNA localization in other neuronal systems, we expressed our reporter transcripts in iPS-derived human motor neurons. As with the CAD and N2A systems, we site-specifically integrated reporter genes into iPS genomes using *cre*/loxP-mediated recombination. Reporter and control iPS cells were then differentiated into motor neurons and mechanically fractionated into cell body and neurite fractions as described previously (Hudish et al., 2020). (**Figure 1A, 5F**). Reporter transcripts containing the mouse *Net1* peak oligonucleotide were 30% more enriched in neurites than a control reporter transcript (p = 0.03, t-test) (**Figure 5G**). This result, combined with the conservation of A/G rich sequences in the 3′ UTRs of the human orthologs of the genes tested in the MPRA (**Figure 4L**), suggests that A/G rich sequences also regulate RNA localization in human neurons.

### Unkempt is required for efficient localization of peak oligonucleotides

In order to identify RBPs required for the efficient transport of our identified peak oligonucleotides derived from *Net1* and *Trak2*, we created biotinylated transcripts containing their sequences and incubated them with CAD and N2A cellular extract. We then retrieved the transcripts from the extract using streptavidin and identified proteins bound to them using mass spectrometry (**Figure 6A**). As a control, we repeated the same procedure using a sequence derived from the open reading frame of firefly luciferase. This sequence was present in all tested reporter constructs, both localized and unlocalized, as well as control reporter constructs (**Figure 5B-E**). It therefore has little intrinsic localization ability and is well-suited to serve as a control in this experiment. Proteins that were specifically bound to the peak oligonucleotides were identified as those more abundant in the peak oligonucleotide RNA pulldown compared to the control RNA pulldown.

**Figure 6.**
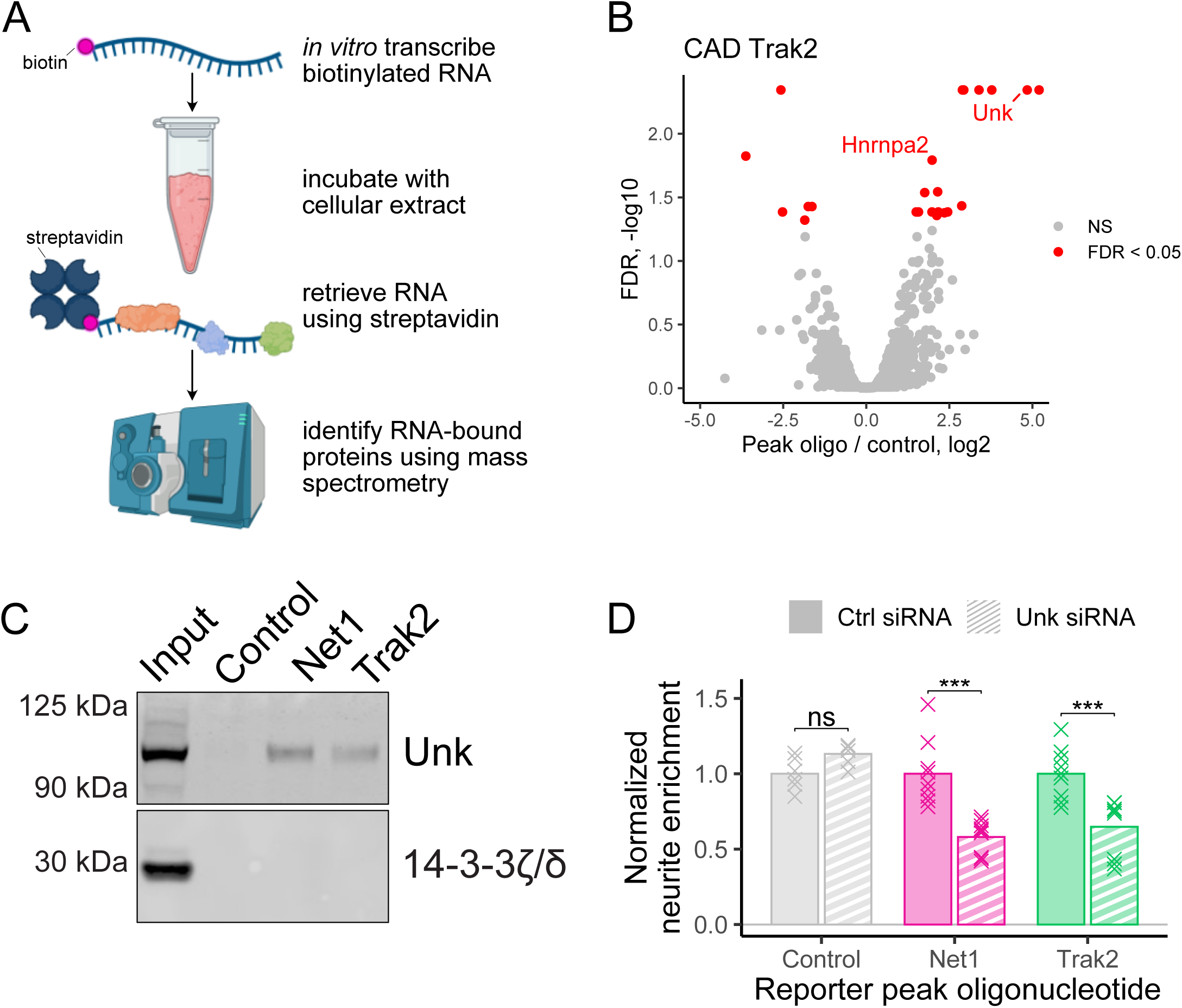
Identification and functional validation of RBPs required for peak oligonucleotide localization. (A) Scheme of mass-spectrometry-based experiment to identify RBPs bound to peak oligonucleotide sequences. (B) RBPs derived from CAD extract that were significantly different in abundance(FDR < 0.05) in the *Net1* peak oligonucleotide RNA pulldown than the control RNA pulldown. (C) Western blot of RNA pulldowns from N2A cell lysate using RNA baits composed of peak oligonucleotides (Net1 and Trak2) or a portion of the coding sequence of firefly luciferase (control). (D) Neurite-enrichments, as determined by cell fractionation and RT-qPCR, of *Net1* and *Trak2* peak oligonucleotide reporter transcripts following the siRNA-mediated knockdown of Unk. All significance tests were performed using a Wilcoxon rank-sum test. p value notation: * < 0.05, ** < 0.01, *** < 0.001, **** < 0.0001.

The only two RNA binding proteins significantly enriched (FDR < 0.05) on both the *Net1* and *Trak2* peak oligonucleotide transcripts in both the CAD and N2A samples were Hnrnpa2 and Unkempt (Unk) (**Figure 6B, S6A-C, Tables S5-S8**). We further confirmed the association of Unk with the *Net1* and *Trak2* probes using immunoblotting (**Figure 6C, S6D**). To assay the ability of Hnrnpa2 and Unk to functionally regulate RNA localization through sequences contained within the *Net1* and *Trak2* peak oligonucleotides, we measured the neurite-enrichment of reporter constructs containing the peak oligonucleotides following Hnrnpa2 and Unk knockdown using siRNAs (**Figure S6E-F**). While the knockdown of Hnrnpa2 expression did not result in reduced neurite localization of either peak oligonucleotide construct (**Figure S6G**), the knockdown of Unk resulted in significant decreases in neurite localization for both constructs (**Figure 6D**). Unk protein is therefore both associated with *Net1* and *Trak2* peak oligonucleotide RNA sequences and required for the efficient neurite localization of transcripts that contain them.

## DISCUSSION

In this study, we identified RNA sequences that were both necessary and sufficient for the transport of a given transcript to neurites. Because the oligonucleotides we used were quite long (260 nt), this allowed us to capture regulatory elements that may also be large. In contrast to short (4-8 nt) RNA elements that regulate processes like alternative splicing and RNA stability, many of the previously identified localization regulatory elements are tens to hundreds of nucleotides long (Engel et al., 2020). The elements that we identified likely require similarly long lengths in order to be fully active. For two of the 260 nt active oligonucleotides that we identified, core subsequences of length 56 and 107 nt drawn from those oligonucleotides had significantly less localization activity than the full length oligonucleotide. The full activity of the element therefore requires more sequence than was contained within the subsequences. This implies that there may be more to the character of these elements beyond simple linear sequence recognition.

A predominant feature of our identified regulatory elements was their peculiar nucleotide composition. Relative to non-enriched oligonucleotides, neurite-enriched oligonucleotides were strongly enriched for adenosine and guanosine residues. We could not discern a more fine-grained sequence enrichment beyond a general preference for adenosine and guanosine. Interestingly, A/G rich sequences have been previously identified in the regulation of localization to projections of mesenchymal cells (Moissoglu et al., 2020), suggesting that such sequences may be general targeting elements for cellular extensions.

RNA localized to the projections of mesenchymal cells through the action of A/G rich sequences can do so through the action of a protein called Adenomatous polyposis coli (APC) (Mili et al., 2008; Moissoglu et al., 2020). In fact, APC was found to bind A/G rich sequences *in vitro* and in cells, lending support to this result (Preitner et al., 2014). We therefore speculated that APC might be involved in the transport of the sequence elements we identified. However, we did not identify APC in our RNA pulldown mass spectrometry experiment. Further, the 3′ UTRs containing our identified localization elements were not bound by APC in a published CLIP-seq dataset (Preitner et al., 2014). Nevertheless, to explicitly test the requirement of APC for the localization of our peak oligonucleotide sequences, we knocked down APC expression using siRNA and monitored the localization of reporter transcripts containing peak oligonucleotides. APC loss did not result in a reduction of localization for the reporters, suggesting that it is not functionally involved with their localization in this neuronal cell type (**Figure S6H**).

The observation of Unk as an RNA binding protein important for efficient localization of peak oligonucleotide-containing transcripts is supported by the involvement of Unk in the maintenance of neuronal morphology (Murn et al., 2015). Interestingly, depletion of Unk results in the loss of neuronal shape while the ectopic overexpression of Unk results in the formation of projections in cells that do not normally have them. The relationship of the regulation of RNA localization by Unk to these functions remains to be seen.

At the time this study was performed, two other groups undertook similar approaches to identify RNA localization regulatory sequences in neuronal cells (Mikl et al., 2021) (Marina Chekulaeva and Igor Ulitsky labs, in preparation). The results of this study are complementary to the findings of these two groups and highlight the power of massively parallel approaches to understand RNA localization mechanisms.

In sum, here we have identified active RNA elements that regulate transcript localization in both mouse and human neuronal cells using an unbiased MPRA approach. Given that hundreds to thousands of RNAs are known to be asymmetrically localized in a variety of cell types, the broad application of MPRAs to RNA localization may be successful in uncovering more elements that govern this process. Further, targeted MPRAs that interrogate specific features within identified elements through mutations and truncations may be useful to determine additional and/or more precise mechanisms that govern RNA localization.

## ACKNOWLEDGEMENTS

We thank members of the Taliaferro lab for helpful discussions regarding experiments and analyses. Portions of the figures in this manuscript were created using Biorender.

This work was funded by the National Institutes of Health (R35-GM133885 to JMT), a Webb-Waring Early Career Investigator Award from the Boettcher Foundation (AWD-182937 to JMT) and the RNA Bioscience Initiative at the University of Colorado Anschutz Medical Campus (JMT). This work was supported by a Culshaw Junior Investigator Award in Diabetes (HAR), the Children’s Diabetes Foundation (HAR). RCG was supported by a CU Anschutz RNA Bioscience Initiative Scholar award.

## METHODS

### Cell culture and subcellular mechanical fractionation

CAD cells were grown in DMEM/F-12 (Gibco, # 11320-033) supplemented with 10% Equalfetal (Atlas Biologicals, #EF-0500-A) and 1% Penicillin-Streptomycin solution. N2A cells were grown in DMEM (Gibco, #11965-092) supplemented with 10% Equafetal (Atlas Biologicals, # EF-0500-A) and 1% Penicillin-Streptomycin solution. The cells were grown in a humidified incubator at 37°C and 5% CO_2_.

For subcellular fractionation, the cells were grown on porous, transwell membranes with a pore size of 1.0 micron (Corning, #353102). Prior to seeding the cells, the bottom of the membrane was coated with 0.2% Matrigel (Corning #356237, diluted in growth medium). Matrigel was allowed to polymerize for an hour at 37°C and 5% CO2. One million cells were plated on each membrane and membrane was placed in one well of a deep six well plate (Corning, # 353502). One 6-well plate constituted one replicate. The cells were allowed to attach for an hour, and then induced to differentiate into a more neuronal morphology. To induce differentiation of the cell lines, medium was changed to differentiation medium (DMEM/F-12 with 1% Penicillin-Streptomycin solution and DMEM with 1% Penicillin Streptomycin solution, for CADs and N2As respectively) and cells were allowed to grow for 48 hours in differentiation medium.

After 48 hours the soma and neurite fractions were harvested as follows, the medium was aspirated, and the membranes were washed 1X with 1 ml PBS. 2 ml PBS was added to the well and 1 ml PBS was added to the top (soma) side of the membrane. The top of the membrane was then scraped gently (to avoid puncturing the membrane) but thoroughly with a cell scraper and the soma fraction were collected into a 15 mL falcon tube on ice. After scraping, the membranes were removed from the insert using a razor blade and were soaked in 500 μL RNA lysis buffer (Zymo, #R1050) at room temperature for 15 min. This constituted the neurite fraction. In parallel, the 6 ml of soma suspension in PBS was spun down at 2000g at 4°C and resuspended in 600 μL PBS. 100 μL of this soma sample was then used for RNA isolation using the Zymo QuickRNA MicroPrep kit (Zymo Research, #R1050). RNA was also isolated from the 500 μL neurite lysate using the same kit. The efficiency of the fractionation was analyzed by Western blotting using ß-actin (Soma and neurite marker, Sigma #A5441, 1:5000 dilution) and Histone H3 (Soma marker, Abcam #ab10799, 1:10000 dilution) antibodies.

### RT-qPCR quantification of reporter transcripts

Two wells of the 6-well plate served as one replicate for the RT-qPCR experiment such that one 6-well plate is seeded per reporter construct to be tested. The cells are plated on the transwell membrane and fractionated as mentioned above. The RNA from both soma and neurite fractions is collected and purified using Zymo QuickRNA MicroPrep kit (Zymo Research, #R1050). cDNA from 100 ng of RNA from each fractions is synthesized using iScript™ Reverse Transcription Supermix (Bio-Rad, #1708841) in a 10 μL volume with a longer incubation time of 30 mins for reverse transcription step. The cDNA is diluted to 20 μL with RNAse-free water. 2 μL of diluted cDNA is used as the template for qPCR to estimate the abundance of Firefly and Renilla transcripts in soma and neurite fractions. The qPCR reaction was performed using Taqman Fast Advanced Master Mix (Life Technologies) with differently labeled probe sets (Life Technologies) allowing to use renilla counts as an internal control. Firefly luciferase RNA was quantified using a VIC-labeled probe while Renilla luciferase RNA was quantified using a FAM-labeled probe.

### Integration of plasmids into cultured cells

CAD and N2A LoxP cell lines were plated in 5X 6-well plates at 5×10^5^ cells per well in respective growth medium 12 to 18 hr before transfection. Cells were then co-transfected with the cloned reporter plasmid mixed with 1% of plasmid expressing Cre recombinase. To transfect one well of a 6-well plate, 1.5 μg of reporter plasmid and 15 ng of Cre-plasmid was mixed with 3 μL Lipofectamine LTX reagent (Invitrogen, #15338100), 1.5 μL PLUS reagent and 100 μL Opti-MEM following the manufacturer’s protocol. Cells were incubated with the transfection mixtures for 24 hours, followed by the media change. The cells were incubated for an additional 24 hr allowing for recovery and expression of the antibiotic resistance before addition of puromycin (2.5 μg/ml). The cells were selected in the puromycin until the cells in the control wells died. The cells with stably integrated reporter plasmids were expanded in the growth medium with puromycin. Three-fourth of the expanded cell lines were frozen, and the remaining were induced with 1 μg/mL doxycycline for 48 hours to express the reporters. After induction, the cells were lysed with TRIzol reagent (Invitrogen, #15596018) and RNA was purified according to the manufacturer’s instructions to analyze the number of integrants and diversity of the library.

### smFISH visualization of reporter transcripts

CAD cells with integrated reporter constructs/peak oligos were seeded on PDL coated glass coverslips (neuVitro, #H-18-1.5-pdl) placed in the 12-well plate at approximately 5 × 10^4^ cells per well. Cells were allowed to attach for 1 hour, followed by media change to serum depleted media with 1 μg/mL doxycycline. Cells were allowed to express constructs and differentiate for 48 hours.

After 48 hours, the media was aspirated, and cells were washed 1X with PBS. Cells were fixed in 3.7% formaldehyde in 1X PBS for 15 min at room temperature, then washed 2X with 500 μL PBS. Next, cells were permeabilized with 70% ethanol at room temperature for 4 hours or at 4°C overnight. The cells were washed with smFISH Wash buffer A at room temperature for 5 min. In the meantime, hybridization reaction was prepared as follows: 2 μL of Stellaris FISH Probe (against Firefly luciferase CDS conjugated with Quasar 670) were added to 200 μL of Hybridization buffer. A hybridization chamber was prepared using an empty opaque tip box with wet paper towels below and parafilm covering the top. Hybridization reaction prepared as above (200 μL per reporter construct) was added on top of the parafilm and the PDL coated glass coverslips were placed with cell side down facing the hybridization reaction. The hybridization chamber with the coverslips was incubated at 37°C overnight. Next day, glass coverslips were transferred to fresh 12-well plates with cell side up and incubated with smFISH Wash Buffer A at 37°C in the dark for 30 min. Samples were stained with fresh smFISH Wash Buffer A supplemented with 100 ng/mL DAPI (Sigma, #D9542-1mg) and incubated in the dark at 37°C for 30 min. DAPI staining buffer was washed with fresh smFISH Wash Buffer B and incubated for 5 min at room temperature. Coverslips were then mounted upside-down onto slides with 6 μL Fluoromount G (Southern Biotech) and sealed with nail polish. Slides were imaged on a widefield DeltaVision Microscope (GE) using 60X objective with 1.4 numerical aperture with 1.33 refractive index oil. The excitation and emission wavelengths used were 547 nm and 583 nm respectively.

### Oligonucleotide library design and synthesis

The code for designing the MPRA oligos is available at https://github.com/TaliaferroLab/OligoPools/blob/master/makeoligopools/OligoPools_shortstep_260nt.py. This script designed 260 nt oligonucleotides with a step size in between neighboring oligonucleotides of 4 nt. For a given gene, oligonucleotides were designed against the 3′ UTRs of all protein-coding transcripts with well-defined 3′ ends (as defined as not having the ‘cds_end_NF’ or ‘mRNA_end_NF’ tags). The polyA site of the UTR was also required to be positionally conserved in humans. This was assessed by getting the syntenic region surrounding the mouse polyA site in the human genome using UCSC liftOver. A polyA site was defined as conserved if there was a polyA site within 200 nt of the corresponding region of the human genome. Additionally, UTRs longer than 10kb were excluded.

UTRs of multiple filter-passing transcripts for a single gene were merged together to create a meta-UTR. Oligonucleotides were then designed against this meta-UTR with the addition of 260 nt upstream and downstream of the beginning and end of the UTR in order to give full coverage of the ends of the UTR with multiple oligonucleotides. 20 nt PCR handles were then added to the ends of every oligonucleotide. The pool of oligonucleotides was synthesized by Twist Biosciences.

### Oligonucleotide library cloning

The oligo pool obtained from Twist Bioscience was resuspended in 10 mM Tris-EDTA buffer, pH 8.0 to a concentration of 10 ng/μl. The libraries for each reporter (Firefly and GFP) were amplified by performing 2X PCR reactions 50 μl each, using Kapa HiFi HotStart DNA Polymerase (Kapa Biosystems, #KK2601) according to the manufacturer’s instructions. We used 10 ng of the original pool as the input DNA template in the reaction and performed 15 cycles in total. The oligonucleotide pool was amplified using primers specific to the 20-nt common sequence and an overhang containing sequence specific to the cloning site for each reporter. After amplification, the PCR reaction was digested with Exonuclease I, at 37°C for 2 hours to digest the single-stranded template and primers. The DNA was then purified using Zymo DNA Clean & Concentrator kit (Zymo Research, #D4013) according to the manufacturer’s protocol.

The reporter plasmid was linearized by digesting it with PmeI (NEB) and BstXI (NEB) at 37°C for 4 hours to clone the library into the 3’-UTR of Firefly and GFP reporter respectively. Digested plasmid DNA was gel extracted using Zymoclean Gel DNA Recovery Kit (Zymo Research, #D4008). The digested plasmid and amplified DNA library were assembled using Gibson Assembly reaction (NEB) using the insert: vector molar ratio of 6:1 at 50°C for 30 mins. The cloned reporter plasmid (∼ 200ng DNA) was purified to get rid of excess salts and was then transformed into E.coli using MegaX DH10B T1R Electrocompetent Cells (ThermoFisher, #C640003), using Biorad GenePulser electroporator. The transformed cells are grown in recovery medium at 37°C for an hour and then plated on pre-warmed Luria broth (LB) agar-Carbenicillin 15-cm plates and incubated at 37°C overnight. The next day, the colonies were collected in a culture tube on ice by scraping the plates using LB medium and spreader. The bacterial culture was pelleted at 4,000 rpm for 20 min. The reporter plasmids libraries were purified using ZymoPURE Plasmid Maxiprep kit (Zymo Research, #D4203). Restriction digestion was performed to confirm that the plasmid library contains only a single insert of the right size.

### Simulation of MPRA quantification

In order to determine if short read aligners like Bowtie2 would be able to accurately assign reads to a set of oligonucleotides that differ from each other by only 8 nt (4 nt on each end), we performed a simulation of the MPRA results using different step sizes. The code for this simulation is available at https://github.com/TaliaferroLab/OligoPools/tree/master/ScreenSimulations.

Oligonucleotides were synthesized from simulated UTRs drawn from the sequence of mouse chromosome 1. UTRs were given random sizes between 500 and 5000 nt. Each oligo was then assigned a random abundance within the simulated MPRA sample. A fastq file of 10 million reads using these abundances was then created. The paired-end fastq files contained the first 97 nt of the oligonucleotide in the forward read and the last 91 nt of the oligo in the reverse read, mimicking the situation encountered in real data after trimming adapters. Deletions and mutations were modeled into the reads at per-base rates of 0.0001 and 0.002, respectively.

This simulated library was then aligned to the reference oligonucleotides using bowtie2 (Langmead and Salzberg, 2012) and the following command: bowtie2 -q –end-to-end –fr –no-discordant –no-unal -p 4 -x Bowtie2Index/index −1 forreads.fastq −2 revreads.fastq -S sample.sam

Quantifications of oligonucleotides from the aligned reads were then compared to the known quantifications produced in the simulation. This process was then repeated with the addition of the bowtie2 parameter -D 150.

### Targeted RNA sequencing of MPRA library

500 ng total RNA from each soma and neurite fractions was taken to synthesized cDNA in a 20 μL reaction using SuperScript IV Reverse Transcriptase (ThermoFisher) according to the manufacturer’s protocol, with primers specific to Firefly and GFP CDS containing an 8-nt unique molecular identifier (UMI) and a partial Illumina read 1 primer sequence. The incubation time at extension step (55°C) was increased to an hour. Post reverse transcription, 1ul each of RNAse H and RNaseA/T1 mix was added directly into the RT-reaction and incubated at 37°C for 30 mins to digest the remaining RNA and RNA:DNA hybrids. The cDNA was purified using Zymo DNA Clean & Concentrator kit (Zymo, #D4013) using 7:1 excess of DNA binding buffer recommended for binding ssDNA.

For library preparation, each purified reporter cDNA reaction was split into 5X PCR reactions (4 μL cDNA/ PCR) and amplified using a reporter specific forward primer with illumina sequencing adaptors and a reverse primer binding the partial illumina read 1 sequence with the remaining sequencing adaptors using Kapa HiFi HotStart DNA Polymerase (Kapa Biosystems, #KK2601) using 18X cycles for GFP and 23X for Firefly reporter. The 5 PCR reactions/sample were pooled together and purified using double SPRI beads protocol, i.e. first purification to get rid of longer non-specific DNA using 0.5X SPRI beads and using the supernatant from the first purification (0.8-0.5 =0.3X) to get shorter non-specific products. The library was quantified using Qubit dsDNA HS Assay Kits (ThermoFisher, # Q32854) and size of the library was verified using Tapestation (Agilent D1000 ScreenTape).

### Alignment and quantification of MPRA results

Adapter sequences were removed from reads using cutadapt (Martin, 2011). Specifically, the sequences GGCGGAAAGATCGCCGTGTAAGTTTGCTTCGATATCCGCATGCTA and CTGATCAGCGGGTTTCACTAGTGCGACCGCAAGAG were trimmed from the 5′ ends of the forward and reverse reads, respectively. The trimmed reads were then aligned to the reference oligonucleotide sequences using bowtie2 and the parameters outlined above (including -D 150). Typically, 99% of reads had the expected adapters, and 95% of those aligned to the reference oligonucleotides.

The number of unique UMIs (the first 8 nt of the reverse read) for each reference oligonucleotide was then calculated using https://github.com/TaliaferroLab/OligoPools/blob/master/analyzeresults/UMIsperOligo.py. These UMI counts were then given to DESeq2 (Love et al., 2014) to quantify oligonucleotide abundances in each sample and identify oligonucleotides enriched in soma or neurite samples.

### Generation, culture and motor neuron differentiation of induced human pluripotent stem cells (hiPSC)

Derivation of induced pluripotent stem cells (iPSC) from a healthy donor was described previously (Taylor et al., 2018). hiPSC were grown as previously described (Hudish et. al, 2020). Briefly, hiPSC were maintained in mTESR plus (Stem Cell Technology, #100-0276) medium on hESC-qualified Matrigel (Corning, #354277) coated cell culture plates. hiPSC exhibited a normal karyotype and were regularly tested for mycoplasma contamination and found negative. Direct differentiation was performed essentially as described previously (Hudish et al., 2020). Briefly, cluster formation was initiated when iPSC cultures reached 70%-90% confluency using microwells (Aggrewells800, Stem Cell Technology) at 3,000 cells per cluster in the presence of 10 uM Rock Inhibitor Y-27632 (RI, Tocris #1254). 24 hours later, clusters were transferred to suspension culture plates at ∼600 clusters per 6-well, maintained on an orbital shaker platform (set at rotational speed 100) in a regular cell culture incubator and motor neuron differentiation was initiated. Mature motor neurons were harvested between days 18-26 for downstream analysis.

### Genetic engineering of hiPSC

hPSC were dissociated into single cells using TrypLE by incubation at 37°C for 8 min. Cells were quenched with mTESR+ media, diluted in PBS, and counted using MoxiGo II cell counter (Orflow). 2×10^6^ cells were transferred into microcentrifuge tubes, spun down and washed twice with PBS. Cells were then prepared for nucleofection of TALEN mediated knock-in (KI) (i) or Cre mediated RCME (ii). Nucleofection was performed using P3 buffer following the Amaxa P3 Primary cell 4D-Nucleofector kit protocol (V4XP-3024) using program: CB-150. (i) 0.5 μg of AAVS1-TALEN-L and AAVS1-TALEN-R (gift from Dr. Danwei Huangfu, Addgene plasmid # 59025) as well as 2 μg of targeting plasmid were used for nucleofection of hPSC cells. Nucleofected hPSC were plated in 10cm plates with 10uM RI, 1x CloneR (STEMCELL # 05888) and 1 μM SCR7 (Excess Bioscience #M60082). 48 hours after plating, blasticidin selection (10 μg/ml) was performed for 10 days. Surviving clonal colonies were picked and expanded for characterization. (ii) For RMCE, 7×10^6^ cells were electroporated with a constitutive Cre plasmid and RIPE cassette using a BioRad GenePulser electroporation system using an exponential decay with 250V and 500 μF settings conditions. Electroporated cells were plated in 10 cm plates with 10 μM RI, and 1x CloneR. 48 hours after plating, puromycin selection (0.5 μg/ml) was performed for 48 hours. Clonal colonies were picked after ∼10 days and expanded for further characterization.

### Computational predictions of RNA structure

RNA structure predictions were performed using RNAfold (Lorenz et al., 2011). For the prediction of guanosine residues participating in quadruplex interactions, the -g flag was used.

### Analyses of sequence conservation

Sequence conservation was assessed using phastCons scores (Siepel et al., 2005) downloaded from UCSC.

### RNA pulldowns

The pull-down experiments and subsequent mass spectrometry were performed according to an adapted experimental pipeline that was previously established (Mikl et al., 2021). Specifically, 10 μL (∼20-35 ng/μL) of cleaned-up PCR amplicon (primer sequences are provided in the table below, used with screening pooled library) were used as template of the *in vitro* transcription (HiScribe™ T7 Quick High Yield RNA Synthesis Kit, #E2050S, New England Biolabs), performed at 37°C for 16 hr, followed by DNase I treatment (37°C for 15 min). IVT RNAs were then cleaned-up and concentrated (DNA Clean & Concentrator-5; #D4013, Zymo Research).

3′-desthiobiotin labeling was carried following the manufacturer’s’ guidelines of Pierce™ RNA 3′ End Desthiobiotinylation (ThermoFisher, #20163). Briefly, ∼115 pmol of each RNA was first subjected to fast denaturation in the presence of 25% v/v DMSO (85°C for 4 min) to relax secondary structures and subsequently labelled at 16°C for 16 hr. RNA binding proteins were isolated using Pierce™ Magnetic RNA-Protein Pull-Down Kit (ThermoFisher, #20164). Briefly, 3′-desthiobiotin labelled RNAs were incubated with magnetic streptaividin-coated beads (50 μL of slurry for each RNA probe) for 30 min at room temperature under agitation (600 RPM in a ThermoMixer, Eppendorf). 200 μg of cell lysates (in Pierce IP lysis buffer; #87787, ThermoFisher) derived from fully differentiated CAD or N2a cells was then incubated with 3′-desthiobiotinylated-RNA/streptavidin beads at 4°C for 1 hr under agitation (600 RPM). 20 μL of each eluate was then analyzed by mass spectrometry.

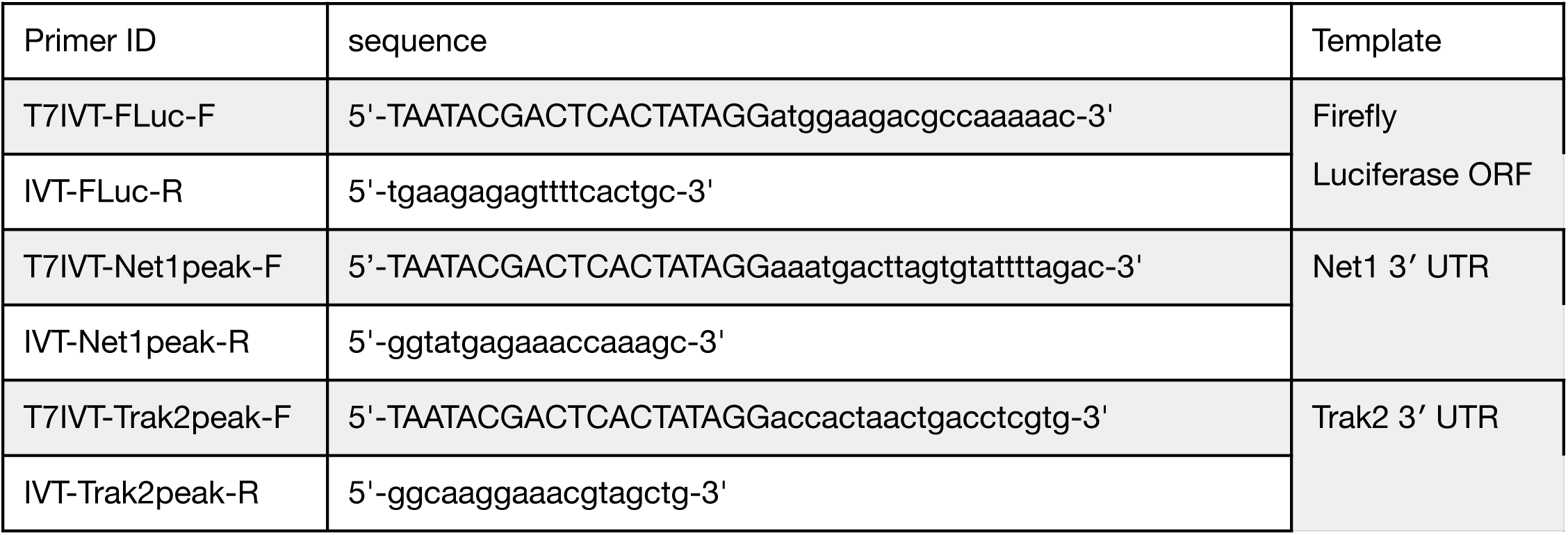

The sequence TAATACGACTCACTATAGG at the beginning of the forward primers encodes a T7 promoter. The biotinylated probe corresponding to the open reading frame of firefly luciferase served as a negative control.

### Mass spectrometry

The pulldown samples were subjected to Trichloroacetic (TCA) precipitation. 20 μL of each eluate was mixed with 80 μL of water and 100 μL of 10% TCA (5% TCA final concentration). The resulting protein pellets were washed twice with cold acetone, dried and dissolved as follows: 45 μl of 10 mM Tris/2 mM CaCl2, pH 8.2 buffer; 5 μl trypsin (100 ng/μl in 10 mM HCl); 0.3 μl trypsin Tris 1M, pH 8.2 to adjusted to pH 8. The samples were then processed with microwave-assisted digestion (60°C for 30 min) and dried. The dried digested samples were dissolved in 20 μL ddH2O + 0.1% formic acid and transferred to autosampler vials for Liquid chromatography-mass spectrometry analysis (LC-MS/MS). 2 μL of sample were injected on a nanoAcquity UPLC coupled to a Q-Exactive mass spectrometer (Thermo Scientific).

The protein identification and quantification was performed using MaxQuant v1.6.2.3 and the data were searched against the Swissprot mouse database.

### RBP knockdowns

Multiple dsiRNAs (3-5 siRNAs/protein) targeting mouse HnrnpA2, Unk and APC were obtained as TriFecta kit from IDT. dsiRNAs were transfected two days prior to differentiation of the CAD reporter constructs in the presence of 1 μg/mL doxycycline followed by 48 hours of differentiation. One 6-well plate per reporter to the tested was transfected using RNAiMax transfection reagent (Invitrogen) according to the manufacturer’s protocol. The efficiency of the knockdown was confirmed by Western blot (HnrnpA2) or qRT-PCR (Unk and APC).

## DATA AVAILABILITY

All high-throughput RNA sequencing data as well as oligonucleotide quantifications have been deposited at the Gene Expression Omnibus under accession number GSE183192.

## SUPPLEMENTARY FIGURES

**Figure S1.**
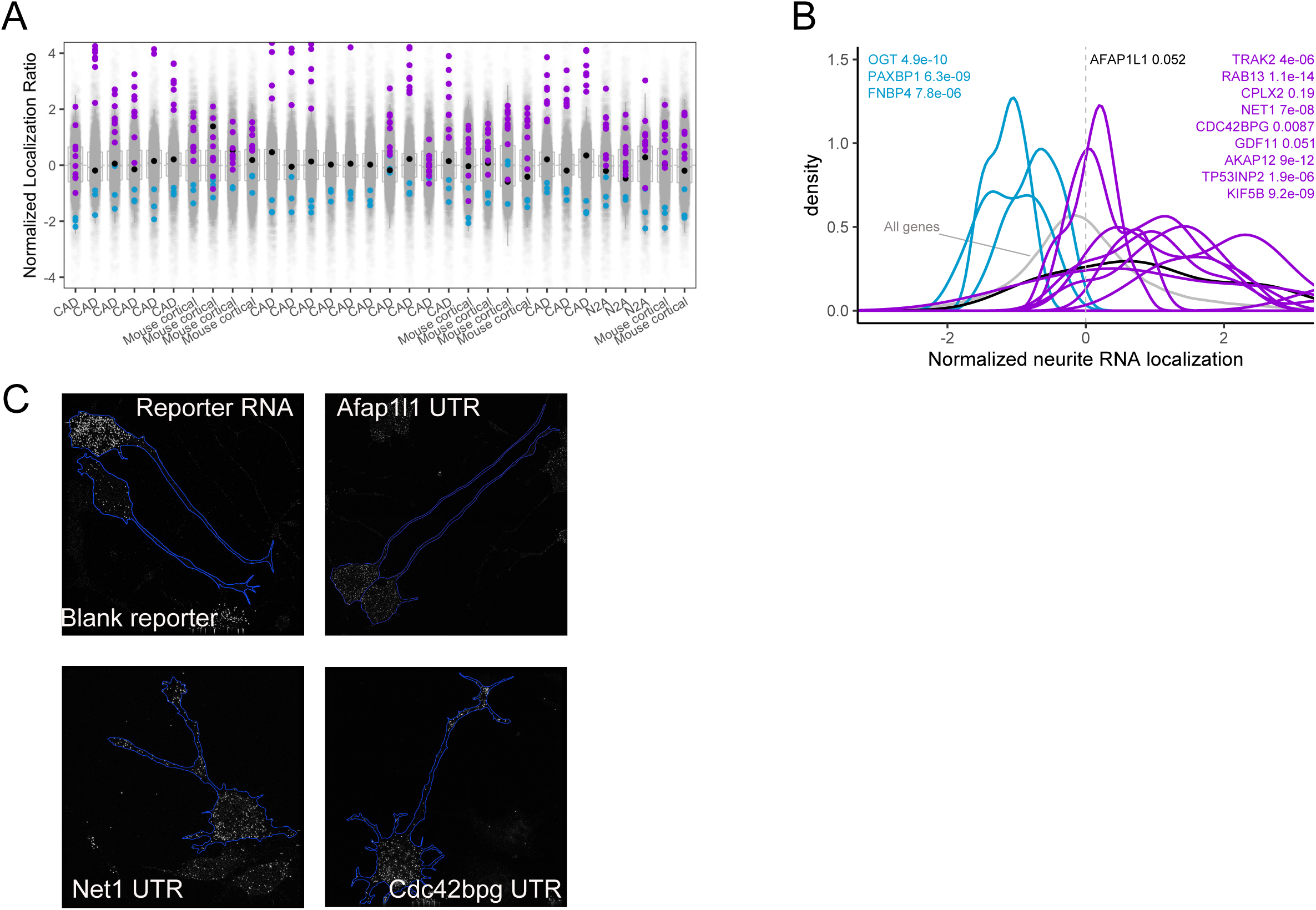
(A) Z-normalized neurite localization values for all genes in all samples quantified. The source of each sample is displayed along the x-axis. Neurite-enriched genes chosen for MPRA analysis are indicated with purple dots. Soma-enriched genes chosen for MPRA analysis are indicated in blue dots. The non-enriched gene chosen for MPRA analysis is in black. (B) iPS-derived human motor neurons were fractionated into soma and neurite fractions. Z-normalized neurite enrichments for the RNAs from selected genes are shown. Genes in purple are orthologs of the neurite-enriched mouse genes chosen for MPRA analysis. Genes in blue are orthologs of the soma-enriched mouse genes chosen for MPRA analysis. The gene in black is the ortholog of the non-enriched gene in the MPRA. Wilcoxon p values represent the differences in neurite localization distributions between the indicated genes and all genes (gray). (C) smFISH imaging of Firefly luciferase reporter transcripts fused to the entire 3′ UTRs of the indicated genes.

**Figure S2.**
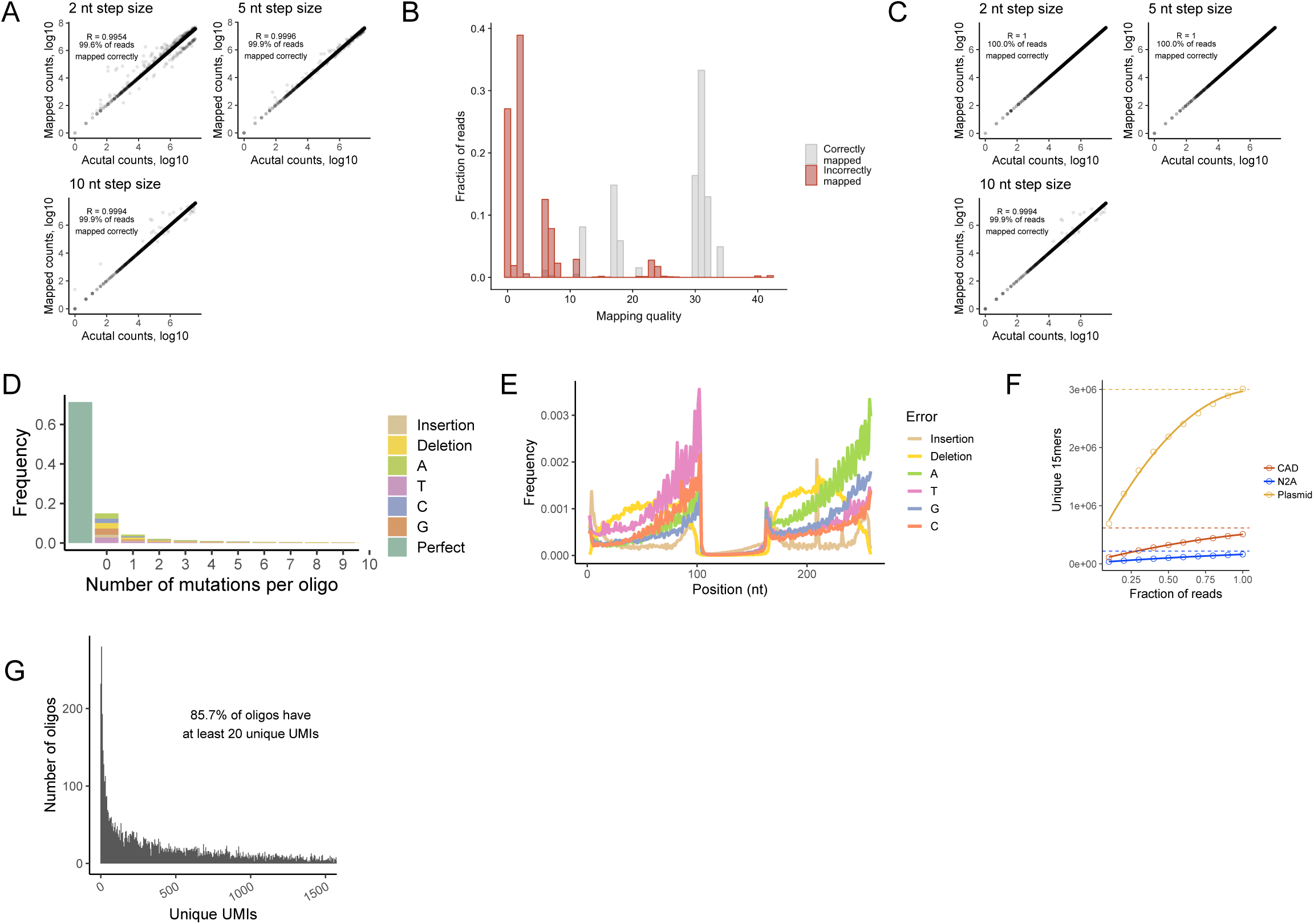
(A) Ten thousand simulated MPRA oligonucleotides were drawn from mouse chromosome 1 using neighboring oligo step sizes of 2, 5, or 10 nucleotides. A mock fastq containing 10 million reads was then made from these oligonucleotides, including 1 nt deletions and mutations at per-base rates of 0.001 and 0.002, respectively. These reads were then aligned to the mouse genome using bowtie2 using the following parameters: bowtie2 -q –end-to-end –fr –no-discordant –no-unal -p 4 -x Bowtie2Index/index −1 forreads.fastq −2 revreads.fastq -S sample.sam The number of reads correctly assigned in each simulated MPRA is shown. (B) Mapping qualities of correctly assigned (gray) and incorrectly assigned (red) reads. (C) Alignment of mock MPRA reads using the same parameters as in (A) with the addition of -D 150. The default value for -D is 15. (D) Number of insertions, deletions, and mutations observed per oligonucleotide in the pool of oligonucleotides.(D) Distribution of error positions across the oligonucleotide. Note that the middle of the oligonucleotide is sequenced by both reads of a paired end sequencing reaction, and errors there were called only if they were present in both reads. The higher rate of errors outside of this middle region therefore likely consists predominantly of sequencing errors. (F) Analysis of the integration efficiency of reporter constructs into cultured cells. Plasmids containing 15mer random sequences were integrated into ∼6 million cells of the integrated line. Following selection for integrants, RNA from each of the line was sequenced, and the number of unique 15mer sequences was calculated. To determine the total number of unique 15mers in the cell population, the number of unique sequences was calculated in subsamples of the data. The relationship between the depth of subsampling and the number of unique 15mers was fit to a quadratic polynomial function, and the number of integrants in the cell pool was defined as the maximum of this function. (G) Distribution of oligonucleotide abundances in the integrated GFP reporter transcript in N2A neuronal cells.

**Figure S3.**
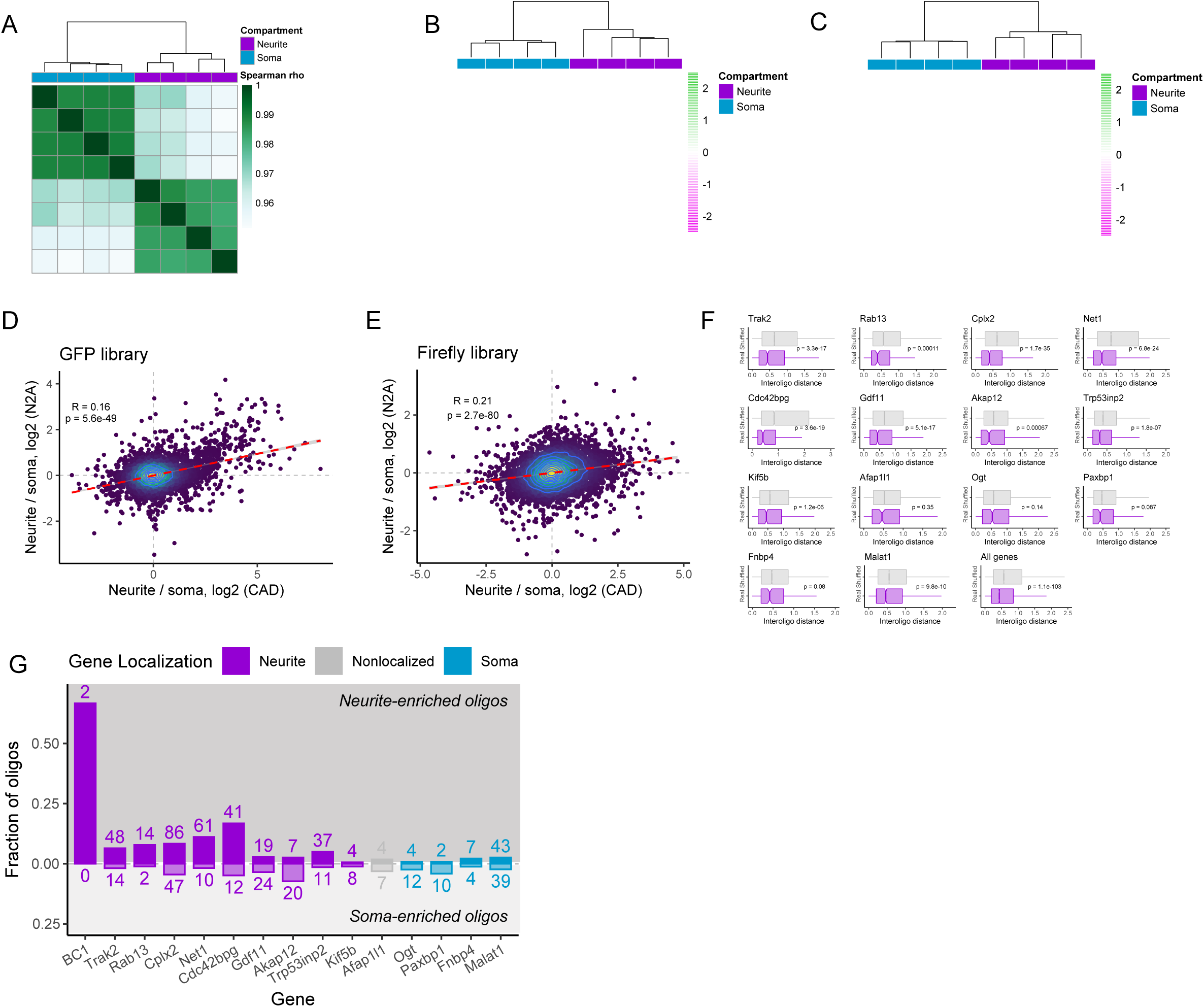
(A) Hierarchical clustering of oligonucleotide abundances from the firefly luciferase reporter in CAD cells. (B) Heatmap of oligonucleotide abundances for all significantly enriched (FDR < 0.01) GFP reporters in the CAD samples. Values are Z-normalized in a row wise fashion. (C) Heatmap of oligonucleotide abundances for all significantly enriched (FDR < 0.01) GFP reporters in the N2A samples. Values are Z-normalized in a row wise fashion. (D) Correlation of neurite enrichments between CAD and N2A samples for all GFP reporters. (E) Correlation of neurite enrichments between CAD and N2A samples for all firefly luciferase reporters. (F) Distribution of absolute differences in neurite enrichment between neighboring oligonucleotides. As a control, the positional relationship between all oligonucleotides was randomly shuffled, and the distances between neighboring oligonucleotides were recalculated. (G) Number of significantly neurite- and soma-enriched oligonucleotides among those drawn from UTRs for each gene.

**Figure S4.**
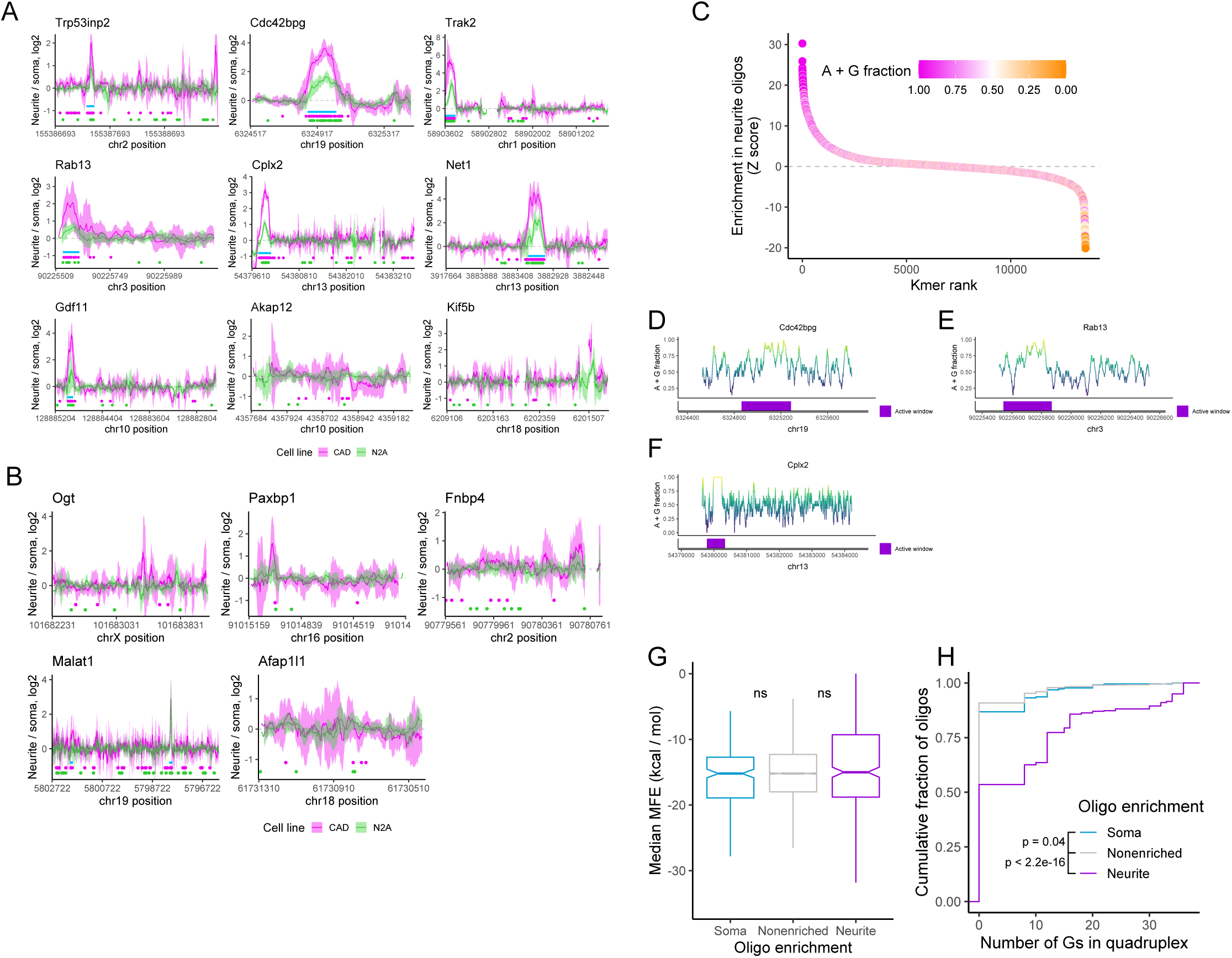
(A) Distribution of neurite enrichment values for oligonucleotides as a function of their position within the 3′ UTR for oligonucleotides taken from the 3′ UTRs of neurite-enriched genes. Data is from the GFP reporter with CAD cell values in pink and N2A values in green. Lines represent a sliding average of 8 oligonucleotides, and the ribbon represents the standard deviation of neurite enrichment for the oligonucleotides in the sliding window. Dots below the lines represent the locations of significantly neurite-localized oligonucleotides (FDR < 0.05). Blue boxes represent the locations of “active windows” defined using the CAD data. Note that the entire 3′ UTRs of *Rab13, Akap12*, and *Kif5b* were not sufficient to drive localization of the reporter transcript. (B) As in A, but for genes that were not neurite-enriched. (C) Kmers enriched in neurite-enriched oligonucleotides as defined by cWords (Rasmussen et al., 2013). (D, E, F) Distribution of A/G content across the 3′ UTRs of *Cdc42bpg* (D), *Rab13* (E), and *Cplx2* (F) and the location of windows of active oligonucleotides, as defined by Figure 4I. (G) The median minimum free energy of oligonucletides defined as soma-, neurite-, or non-enriched as calculated by RNAfold (Lorenz et al., 2011). To compare groups, Wilcoxon ranksum tests were performed. (H) The number of guanosine residues participating in G-quadruplexes of oligonucletides defined as soma-, neurite-, or non-enriched as calculated by RNAfold (Lorenz et al., 2011). To compare groups, Wilcoxon ranksum tests were performed.

**Figure S5.**
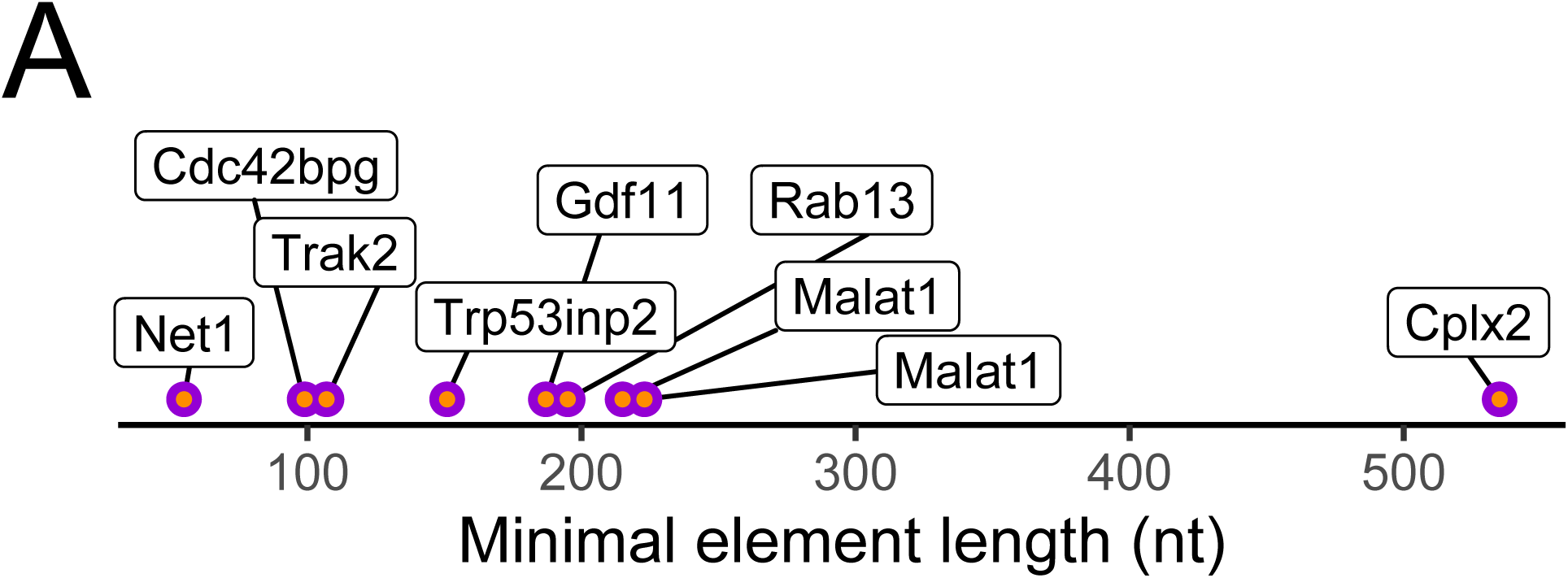
(A) Length of minimal elements, as defined by figure 4I. Cplx2 is an outlier in this analysis as the length of the active window was so long such that there was no sequence present in every active oligo. For this gene, then, the length of the active window is shown.

**Figure S6.**
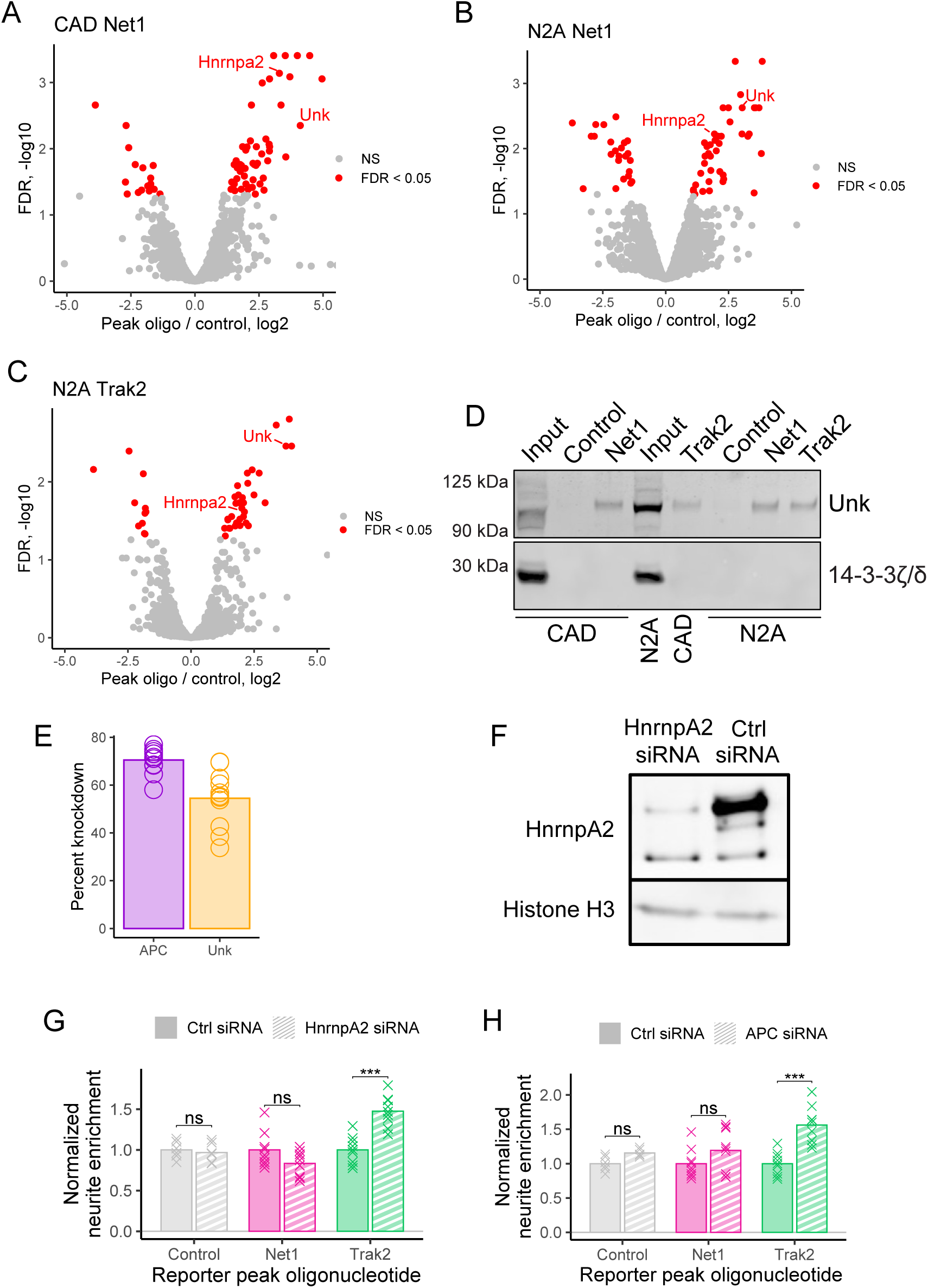
(A) RBPs derived from CAD extract that were significantly different in abundance (FDR < 0.05) in the *Net1* peak oligonucleotide RNA pulldown than the control RNA pulldown. (B) As in A, but using N2A cellular lyaste. (C) As in B, but using the *Trak2* peak oligonucleotide as the RNA bait. (D) Western blot of RNA pulldowns from N2A and CAD cell lysate using RNA baits composed of peak oligonucleotides (Net1 and Trak2) or a portion of the coding sequence of firefly luciferase (control). (E) Efficiency of APC and Unk knockdown as measured by RT-qPCR. (F) Efficiency of Hnrnpa2 knockdown as measured by western blotting. (G) Neurite-enrichments, as determined by cell fractionation and RT-qPCR, of *Net1* and *Trak2* peak oligonucleotide reporter transcripts following the siRNA-mediated knockdown of Hnrnpa2. (H) As in G, but following the knockdown of APC. All significance tests were performed using a Wilcoxon rank-sum test. p value notation: * < 0.05, ** < 0.01, *** < 0.001, **** < 0.0001.

## SUPPLEMENTARY TABLES

**Table S1**. Enrichments of all oligonucleotides for the GFP reporter in CAD cells.

**Table S2**. Enrichments of all oligonucleotides for the Firefly reporter in CAD cells.

**Table S3**. Enrichments of all oligonucleotides for the GFP reporter in N2A cells.

**Table S4**. Enrichments of all oligonucleotides for the Firefly reporter in N2A cells.

**Table S5**. Enrichment of proteins associated with the Net1 peak oligonucleotide RNA as compared to a control RNA sequence in CAD cell extract as determined by mass spectrometry.

**Table S6**. Enrichment of proteins associated with the Trak2 peak oligonucleotide RNA as compared to a control RNA sequence in CAD cell extract as determined by mass spectrometry.

**Table S7**. Enrichment of proteins associated with the Net1 peak oligonucleotide RNA as compared to a control RNA sequence in N2A cell extract as determined by mass spectrometry.

**Table S8**. Enrichment of proteins associated with the Trak2 peak oligonucleotide RNA as compared to a control RNA sequence in N2A cell extract as determined by mass spectrometry.

## SUPPLEMENTARY FILES

**Supplementary file 1**. Sequences for the oligonucleotides used in the MPRA. These sequences do not contain the 20 nt adapters used as PCR handles that were fused to either end of each oligo.

**Supplementary file 2**. Genomic coordinates for each oligonucleotide member of the MPRA.

